# Spinal interneuronal populations encode static hindlimb posture in the cat

**DOI:** 10.1101/2025.11.06.686917

**Authors:** Yuta Soga, Kazutaka Maeda, Shiro Egawa, Enrique Contreras-Hernandez, Shusei Fukuyama, Mirai Takahashi, Kaoru Takakusaki, Tetsuro Funato, Kazuhiko Seki

## Abstract

Proprioceptive signals from primary afferents reflect changes in single-joint angles, whereas neuronal population in the cerebral cortex represent whole-limb postures. Where and how this transformation emerges along the somatosensory axis from peripheral proprioceptive receptors remains unclear. We simultaneously recorded many lumbosacral spinal neurons in two decerebrate, immobilized cats while a robotic device held the hindlimb at 16 static endpoint positions spanning hip–knee configurations. Using high-density multielectrode recordings, we asked how spinal populations encode static limb state. At the single-neuronal level, activities in a majority of neurons covaried with a single joint’s angle (hip or knee), a smaller subset showed combined modulation by both joints, and a distinct subset (‘single-endpoint’ neurons) fired selectively at one unique hip-knee configuration near the sampled joint-range limits and was quiescent in adjacent postures.

Population analyses revealed a low-dimensional structure: the first two principal components tracked knee and hip angles, respectively, whereas a third component isolated a boundary-aligned, posture-specific pattern, with loadings peaking at the same extreme configurations preferred by single-endpoint neurons. Decoders trained on ensemble activity reconstructed both joint angles and the limb’s endpoint position in body-centered coordinates, indicating that the recorded spinal-interneuron populations contain sufficient information to reconstruct whole-limb kinematics. Together, these findings are consistent with a hierarchical organization whereby joint-based representations within spinal-interneuron populations could contribute to the emergence of limb-centered representations in the ascending proprioceptive pathways. The boundary preference of single-endpoint neurons supports a possible categorical coding scheme at workspace limits that may provide spinal “landmarks” for switching control modes, enhancing stability near kinematic extremes, and supporting recalibration of proprioceptive population codes.

**Key Points:**

- We simultaneously recorded many lumbar spinal neurons in two decerebrate, immobilized cats while a robot held the hindlimb at 16 static positions to test how spinal populations encode posture.
- Many neurons varied with a single joint angle (hip or knee), a smaller subset showed combined hip–knee modulation, and a distinct subset was active only at one specific endpoint posture.
- Population analyses revealed a low-dimensional structure: the first two principal components tracked knee and hip angles, while a third captured a posture-specific pattern aligned with single-endpoint neurons.
- Decoders trained on ensemble activity reconstructed joint angles and the limb’s endpoint, indicating that spinal interneuronal populations contain sufficient information for whole-limb kinematics.
- These findings are consistent with a hierarchical organization whereby joint-based representations within spinal-interneuron populations could contribute to the emergence of limb-centered representations in the ascending proprioceptive pathways.

## Introduction

It has been shown that the static postural information about limb is encoded in a graded fashion along the proprioceptive sensory pathway in the nervous system. In studies of single joint- and muscle afferents, firing rates vary linearly with joint angles (Stein *et al*., 2004) (Vallbo, 1974) (Wei *et al*., 1986) with some afferents increasing activity at movement extremes (Clark & Burgess, 1975; Grigg, 1975). At the cortical level, such static configurations of limb are known to be represented in population activity. In nonhuman primates, simultaneously recorded ensembles in primary motor and somatosensory cortices encode multi-joint hand and arm posture during hold epochs; joint-angle combinations (hand “shape” or arm configuration) can be decoded from neural population activity even when movement is absent or minimized (Goodman *et al*., 2019) (Chowdhury *et al*., 2020) (Klaes *et al*., 2015). How joint-based representations at the level of primary afferents are transformed into shape- and configuration-based representations in the cortex, and how single-afferent signals are integrated into population-level representations, remain fundamental questions in neuroscience.

A landmark series of studies by Bosco and Poppele (Bosco & Poppele, 2001) showed that dorsal spinocerebellar tract (DSCT) neurons—an output pathway relaying spinal signals to the cerebellum—encode whole-limb kinematic parameters (Bosco & Poppele, 1993), such as hindlimb endpoint position and leg configuration, rather than merely reproducing receptor- or single-joint–based signals present in primary afferents. In other words, joint-based reference frames (Soechting & Flanders, 1992) in the primary afferents are transformed into limb- or body-centered coordinates at the level of the DSCT.

Building on these works, the present study examines the possibility that such coordinate transformations arise upstream of the spinal ascending pathway within polysynaptic spinal circuits from primary afferent to the neurons ascending to the Brain (e.g., DSCT neurons). We hypothesized that ensemble-level integration occurs within the spinal gray matter, with DSCT neurons relaying, in part, this integrated proprioceptive state to the cerebellum (Jankowska & Puczynska, 2008; Krutki *et al*., 2011). Previously, under anesthesia and spinalization in monkeys, we found that focal excitability of the cervical spinal cord, probed with intraspinal micro-stimulation, changed nonlinearly with passive changes in arm configuration (Yaguchi *et al*., 2015). This observation further supports the hypothesis that multiple proprioceptive afferent inputs, arising from the same or different joints, converge within focal spinal regions that are outside of the DSCT area, providing a potential substrate for coordinate transformation.

Recent primate studies of cortical proprioception (Goodman *et al*., 2019) (Chowdhury *et al*., 2020) (Klaes *et al*., 2015) using simultaneous multi-neuronal recordings have yielded substantial insights into population-level representations across cortical areas. By contrast, most investigation on the spinal proprioceptive representation have relied on single-unit recordings; consequently, prior findings largely describe cell-wise features, and how proprioceptive signals are represented at the level of spinal populations remains unclear. Here, as a step toward closing this gap, we applied high-density multielectrode recordings in the spinal cord and examined how neuronal ensembles respond to systematic changes in hindlimb endpoint position.

## Method

### Animals

All the procedures of the present experiments were approved by the Animal Studies Committee of Asahikawa Medical University (approval number; R4-101) and were following the Guide for the Care and Use of Laboratory Animals (NIH Guide), revised in 1996. Every effort was made to minimize animal suffering and to reduce the number of animals used. The study was based on two adult female cats weighing 2.8 and 3.2 kg, respectively. These cats were obtained from a laboratory animal supplier and bred at the animal facility of Asahikawa Medical University.

### Surgical Operation

In preparation for the measurement, each cat was anesthetized with isoflurane (Viatris Co., Tokyo, Japan; 0.5–3.0%) and nitrous oxide gas (0.5–1.0 l/min) in oxygen (3.0‒5.0 l/min). Under sufficient anesthesia, the trachea was intubated, and a cannula was placed in the right femoral artery to monitor blood pressure. Another cannula was placed in the right radial vein to administer the muscle relaxant (pancuronium bromide, Myoblock, Sankyo Co., Tokyo, Japan) via injection (0.1 mg/kg). A laminectomy of the vertebral arches from T10 to S2 was performed to expose the spinal cord segments. In addition, bipolar electrodes (2–3 mm inter-electrode distance) were attached to the nerve innervating the posterior biceps semitendinosus (PBSt) and the lateral gastrocnemius-soleus (LG-S) muscles on the left side.

The cat was then surgically decerebrated at the precollicular–postmammillary level, and the anesthesia was subsequently discontinued. The head was then fixed in a brain stereotaxic frame, and the cat was secured by a rigid spinal frame clamping the dorsal processes of the first three thoracic vertebrae and the bodies of the first and seven lumbar vertebrae (L1, L7), with both iliac crests additionally secured to the stereotaxic frame using hip pins (Figure 1). The exposed lumbo-sacral spinal cord was covered with a pool of paraffin oil. One hour after the discontinuation of anesthesia, clear decerebrate rigidity appeared, and the cat exhibited a reflexive standing posture accompanied by tonic contraction of the bilateral soleus muscles.

**Figure 1.**
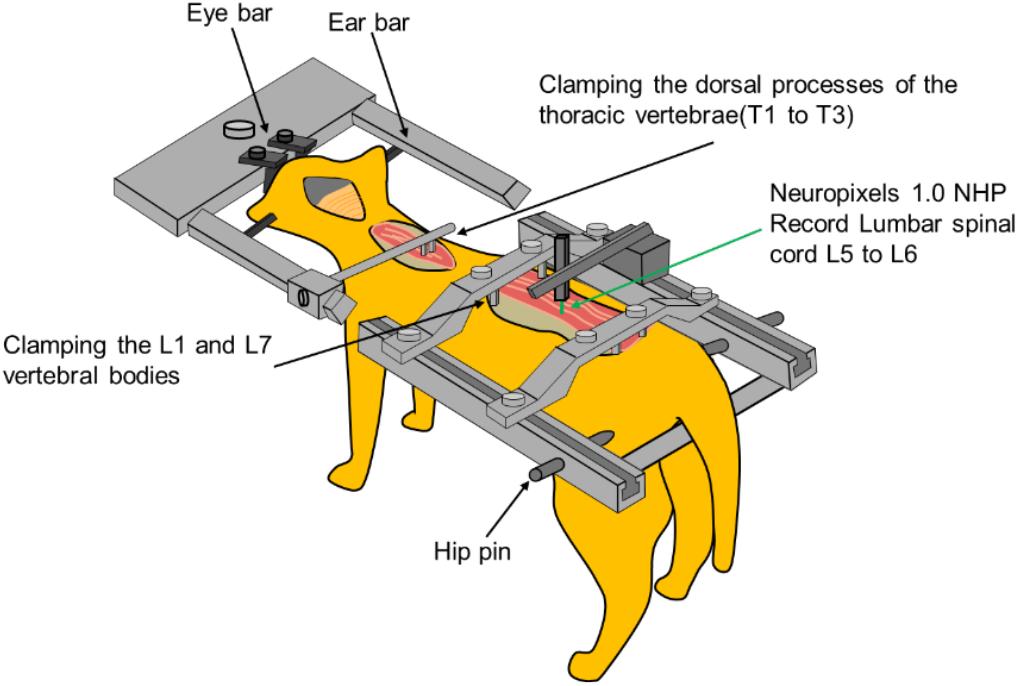
Experimental setup. The decerebrate cat was secured to the stereotaxic frame by fixing the head, the thoracic and lumbar spinal regions, and the iliac crests. The head was fixed to the stereotaxic frame using a brain stereotaxic frame, including the eye bars and ear bars. The dorsal processes of the first three thoracic vertebrae (T1–T3) and the bodies of the first and seven lumbar vertebrae (L1, L7) were clamped. Hip pins were attached to the iliac crests of both hindlimbs. The Neuropixels electrode was attached to the tip of a manipulator mounted on the stereotaxic frame and inserted into the L5/L6 intersegmental space of the left lumbar spinal cord. The animal maintained a reflexive standing posture accompanied by tonic contraction of the bilateral soleus muscles due to decerebrate rigidity.

Prior to the penetration of the recording electrodes, the cat was immobilized by administering a muscle relaxant (Myoblock, 0.1 mg/kg) via the right radial vein. Subsequently, respiration was managed under a respirator to maintain the end tidal CO_2_ concentration between 4 % and 6 %. The mean blood pressure was also maintained above 100 mmHg. Since respiration and blood pressure were highly influenced by body temperature, the temperatures of the animal’s rectum and the oil pool were monitored and maintained at 36–37 °C using radiant heat lamps.

### Measurement of Neural Activity

Electrical stimulation (0.2 ms of rectangular wave at 1.0 Hz with an amplitude 1.2–1.5 times the threshold) was delivered to the nerve fibers innervating the PBSt and the LG-S muscles, respectively. Cord dorsum potentials (CDP) associated with Group I fiber activity evoked by each stimulus were recorded. For recording CDP, a silver ball electrode (diameter 0.5-0.7 mm) placed on the dorsal surface of the seventh lumbar segment L7 was used, and a platinum electrode embedded in the skull served as the reference.

Next, a Neuropixels 1.0 NHP electrode (IMEC, Leuven, Belgium) was fixed to a custom-made manual manipulator (Narisige, Tokyo, Japan) mounted on the stereotaxic frame and inserted from the dorsal to the ventral side of the lumbar spinal cord. The electrode insertion sites were 1 mm lateral to the midline in the sixth lumbar segment L6 and 1 mm lateral to the midline in L5 for Cat A, and 1.2 mm lateral to the midline in L6 for Cat B. At that time, the penetration of the electrode into the ventral grey matter was confirmed by comparing the Group I fiber potential obtained from the CDP with the activity recorded from the inserted electrode. This procedure enabled the detection of neural activity across the grey matter, from the dorsal to the ventral horn. Signals obtained from the Neuropixels probe electrodes were amplified by a headstage (HS_1000, IMEC, Leuven, Belgium), and then acquired via a PXIe Card (PXIE_1000, IMEC, Leuven, Belgium) into the recording software Open Ephys (Siegle *et al*., 2017) (sampling frequency of 30 kHz). The signals were then processed with a high-pass filter, and the spike signals of neural activity were recorded.

### Experiment

The posture of the left hindlimb of the cat secured to the stereotaxic frame was manipulated using a robot arm (DOBOT Magician E6, Shenzhen Yuejiang Technology Co., Ltd., Shenzhen, China). The red markers in Figure 2A show the locations of the kinematic landmarks used for tracking the hindlimb posture. Small beads were attached to the cat’s skin surface to serve as these landmarks. As shown in Figure 2B, a 3D-printed limb interface fixture was attached to the hindlimb at the ankle joint, while the remaining parts of the limb (hip and knee joints) were kept free to move without any external constraints. The limb interface fixture was attached to an acrylic plate fixed to the end-effector of the robot arm, thereby connecting the cat’s hindlimb to the robot. Using the robot arm, the hindlimb was moved to 16 recording positions, varying the angles of the hip and knee joints.

**Figure 2.**
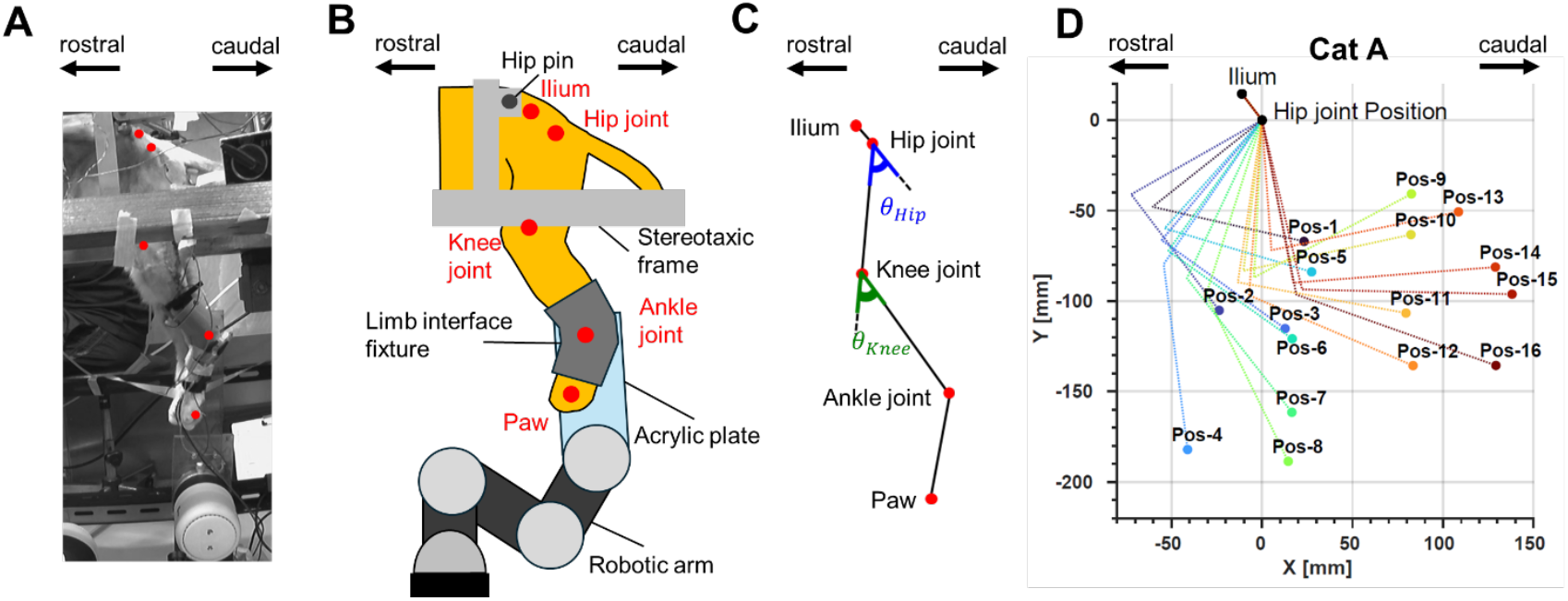
Posture estimation at the recording positions of the cat hindlimb. **A**, The cat hindlimb secured in the stereotaxic frame. Each red marker indicates a landmark for tracking the hindlimb postural information, representing the locations of the ilium, hip joint, knee joint, ankle joint, and paw. These landmarks were the beads attached to the limb interface fixture at the ankle joint and the beads attached to the body surface at the other joints (iliac crest, hip joint, knee joint, and paw). The hindlimb shown in the figure is the posture maintained in a reflexive standing posture during the inter-trial interval. **B**, Connection of the cat hindlimb to the robot. The left hindlimb of the cat, with the iliac crest fixed by a hip pin, was attached via the limb interface fixture to an acrylic plate mounted at the tip of the robotic arm. Each red marker represents the locations of the iliac, hip joint, knee joint, and ankle joint, whose positions were estimated during motion analysis. **C**, Definition of the hip and knee joint angles. *θ*_*Hip*_ denotes the hip joint angle, and *θ*_*Knee*_ denotes the knee joint angle. **D**, XY positions of the ilium, hip, knee, and ankle joints for Cat A estimated by posture analysis. The endpoint represents the position of the ankle joint. Each coordinate point is the time-averaged value of coordinates from 1 to 14 seconds after the start of measurement, averaged across five trials for the 16 recording positions. Note that because the cat’s iliac crest was fixed to the frame by the hip pins, resulting in almost no movement of the iliac crest and hip joint, their coordinates were averaged across all 16 recording positions.

When the hindlimb reached each recording position, it was kept stationary for 15.01±0.01 seconds in both Cat A and Cat B. The hindlimb then moved to the next recording position, with each transition being completed by the robotic arm in less than 2.04 seconds. A 60-s interval was provided between trials, during which the hindlimb was maintained in a reflexive standing posture as shown in Figure 2A. This procedure was repeated until five trials were completed at each position, for a total of eighty trials. To minimize the effects of path dependency and time-dependent changes in neural activity, the order in which the recording positions were visited was randomly varied across trials. To detect and record when the hindlimb became stationary, the robot generated a digital signal upon reaching a recording position and continuously output this signal until the next transition began. This digital signal was fed into the Open Ephys system via a recording board via a terminal block (BNC-2110, National Instruments Corporation, Austin, USA).

### Motion Analysis

To calculate the hip and knee joint angles at each recording position, coordinates for the iliac, hip, knee, and ankle joint locations were estimated from three experimental video recordings. The three experimental videos were recorded at 30 f.p.s. by a camera (WM-F570A, Sony Corporation, Tokyo, Japan) and collected by a recording system (YKS-TN2008AHD-H, YKS Co., Ltd., Osaka, Japan). During the experiment, the three cameras simultaneously recorded a 1Hz flashing LED, and the recorded LED signals were later used for synchronization. 3D DeepLabCut (ver 2.1.6.4, 2.3.11) (Mathis *et al*., 2018) (Nath *et al*., 2019), which enables the estimation of the 3D coordinates of an object simultaneously recorded by two cameras (a camera pair), was used to estimate the coordinates of the iliac, hip, knee, and ankle joint locations.

The 3D position estimation using 3D DeepLabCut was performed as follows. First, referring to the landmarks attached to the surface of the cat’s hindlimb (Figure 2A), only the clearly visible points corresponding to the iliac crest, hip joint, knee joint, and ankle joints were annotated on the 2D images from all three video cameras. Next, a convolutional neural network model (ResNet-50) was trained 800,000 times using the landmarks and their surrounding image regions. For camera-pair calibration, a checkerboard with 10 mm times 10 mm squares was measured from multiple angles, and calibration parameters, including the actual spatial scale, lens distortion, and the relative positions between the cameras, were calculated. Using these calibration parameters together with the trained model, the 3D positions of each landmark were estimated.

The locations of the knee and ankle joints at each recording position were calculated by averaging the coordinate data over five trials, using the data from 1 to 14 seconds after reaching the recording position. Since the iliac crests were fixed to the stereotaxic frame with the hip-pins and thus remained immobile, the positions of the iliac marker and hip joints were determined by calculating the mean coordinate across all 16 recording positions, using data from 1 to 14 seconds after the onset of the stationary period, which served as the representative single coordinate for all postures. In Cat A, the knee joint position could not be estimated at one recording position (Pos-9) because it was obscured by the robotic arm during video recording. Therefore, the knee joint position at Pos-9 was estimated using a two-segment model consisting of the hip–knee and knee–ankle links. The lengths of these two links were set to the average values obtained from all recording positions except Pos-9. The knee joint coordinates at Pos-9 were then computed as the complementary value by applying the ankle position coordinates at Pos-9 to the distal end of the two-link model. Finally, using the estimated coordinates of the iliac crest, hip joint, knee joint, and ankle joint at each recording position, the hip and knee joint angles were calculated according to the definition shown in Figure 2C.

### Pre-processing of Neural Activities 1: Identification of single spike activity using spike sorting

We extracted the spike timings of individual neurons from the signals recorded with the Neuropixels probe and performed offline spike sorting to analyze the activity of each unit. First, we performed electrode drift correction and extracted candidate clusters for single units automatically using Kilosort4 (Pachitariu *et al*., 2024), which performs analysis based on a template matching algorithm. Next, because the automatically classified clusters could include errors, such as noise, over-splitting, or under-merging, we performed manual curation using *Phy*, a Python-based spike-sorting tool. During this process, clusters were excluded, separated, and merged based on waveform shape, cross-correlograms (CCGs), and inter-spike intervals (ISIs). Finally, we evaluated the quality assessment of each cluster following manual curation based on the following two quantitative metrics:

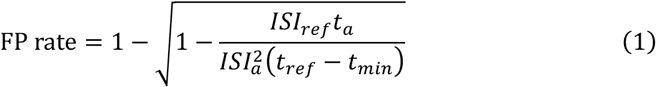

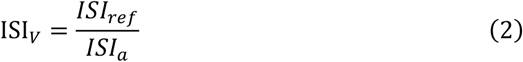

where the FP Rate (false positive rate) in Eq. 1 is a quality metric based on the physiological property of the neuronal refractory period. It indicates the proportion of spikes from other units that are incorrectly included in the target unit (Llobet *et al*., 2022). The refractory period was determined to be 1.5 ms with reference to the 1 ms refractory period reported previously, adopting a more stringent criterion (Prut & Perlmutter, 2003). The parameter *t*_*min*_ is the minimum interval used to exclude artifacts originating from identical waveforms. In Eq. 1, *t*_*a*_ denotes the total recording time, *ISI*_*a*_ denotes the total number of ISIs, and *ISI*_*ref*_ denotes the total number of ISIs falling within the refractory period. Note that *ISI*_*ref*_ that appeared within *t*_*min*_ were excluded as artifacts and not included in the total count. In Eq. 2, ISI_*v*_ denotes the proportion of ISIs included within the refractory period.

We defined a single units as one that satisfies the following criteria for the FP rate in Eq. 1 and the ISI_*v*_ in Eq. 2.

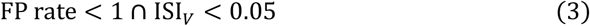

Subsequently, for single units, the entire recording period was divided into five segments, and units that exhibited consistent waveform shape across these segments were retained for further analysis. In this study, axonal and somatic spikes were not distinguished. Also, the data recorded with the electrode inserted into L5 in Cat A were excluded from the analysis because the number of recorded single units was insufficient.

### Pre-processing of Neural Activities 2: Identification of neural activity duration

To examine the activity of single neurons, the spike timing of each neuron was aligned to t=0, which was defined as the moment the cat’s hindlimb achieved the recording position. The spike timing was then converted into firing rates using a 50-ms bin width, and these results were averaged across five trials, as shown in Figure 3A. Subsequently, neuron IDs were assigned sequentially to each neuron, starting from the neuron recorded at the electrode tip. The resulting firing rates for recorded neurons were then visualized as shown in Figure 3B. Some neurons shown in Figure 3B exhibited increased activity during the transition to the next recording position, with a slight residual effect persisting after the t=0 onset. Therefore, to mitigate the influence of this post-effect, we identified the stationary period based on neural activity.

**Figure 3.**
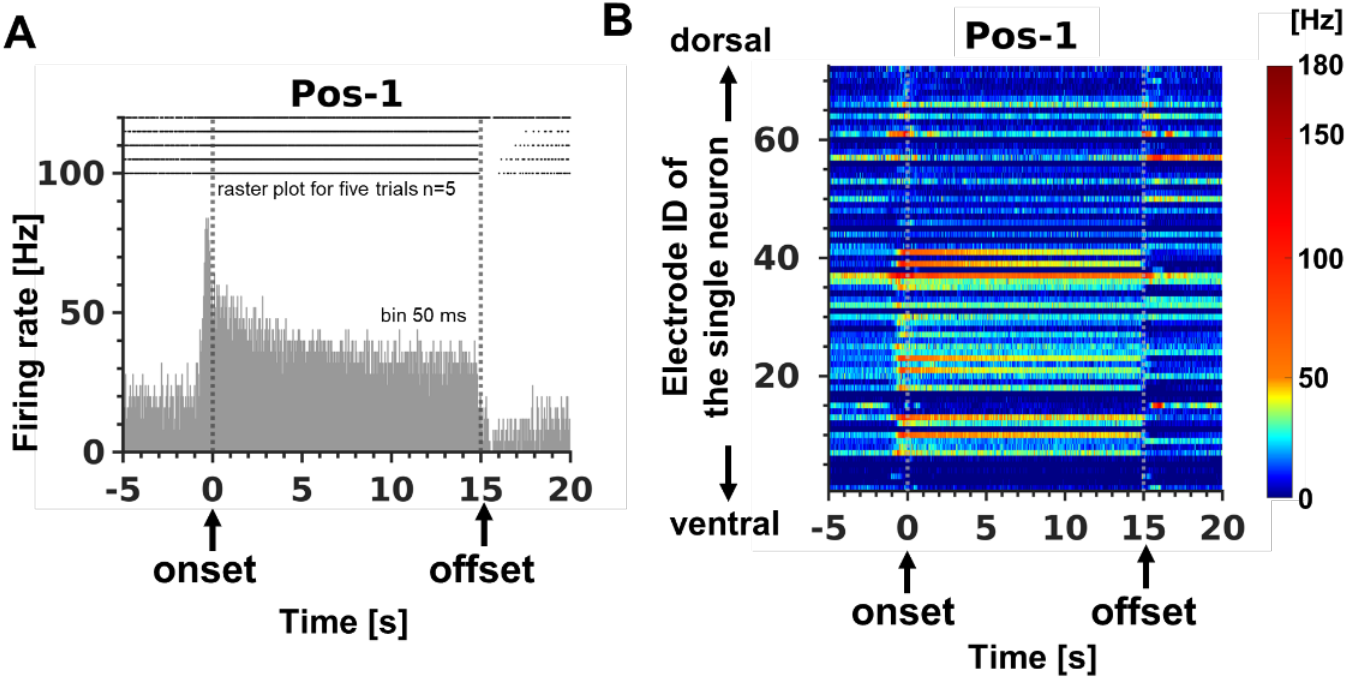
Visualization of recorded neural activity. **A**, Single neuron activity during a particular recording position. The vertical axis of the panel indicates firing frequency, and the horizontal axis indicates time. The dots at the top of the panel indicate the timing of neural activity for a given trial (n=5), and the bar graph represents the firing frequency calculated from the raster plot of five trials using a 50 ms bin width. The dashed line at 0 on the horizontal axis indicates the time of arrival at recording positions (onset of the recording position), and the dashed line at 15 indicates the beginning time of the transition to the next recording position (offset of the recording position). **B**, The activity of all single neurons during a representative recording position. The vertical axis of the panel illustrates the electrode ID of single neurons, with IDs assigned sequentially (1, 2, etc.) starting from the neuron recorded at the electrode tip. These neurons are arranged from ventral to dorsal relative to the electrode tip. The horizontal axis illustrates time. Firing frequency is calculated from the raster plot of five trials using a 50 ms bin width, with its intensity represented by color. The dashed line at time 0 s on the horizontal axis indicates the onset of the recording position, and the dashed line at 15 s indicates the offset. The time intervals represent activity during the pre-recording period(−5 to 0 s), the recording position period (0 to 15 s), and the post-recording period (15 to 20 s).

For this purpose, we first calculated the firing rate of each neuron identified as a single unit for every trial and recording position. Next, we computed the gradient of the averaged firing rate time series (rectified to positive values) using the linear least squares method within a moving window. The moving window spanned nine bins (450 ms), and the gradient for each central bin was calculated using the activity data from the four preceding and four following bins. Based on the calculated firing rate time series, the onset of the stationary period was defined as the first time point at which the gradient changed from negative to positive immediately after reaching a recording position, and the end of the stationary period was defined as the time point where the gradient again showed a large deviation. The criteria for determining the stationary period were calculated using the data from Cat A, where many neurons exhibited a rapid increase in activity immediately after reaching the recording position. The same time range was applied to Cat B because the experimental protocol was identical. For further analysis, the average firing rate of each neuron during this identified stationary period was used as the explanatory variable.

### Relationship Between Single-Unit Activity of Neurons and Joint Angle

To investigate the extent to which the recorded neural activity reflects the hip and knee joint information at each recording position, the relationship between each single-unit activity and the angles of the each joints was analyzed. The analysis was performed using the following multivariate regression model (Bosco *et al*., 1996):

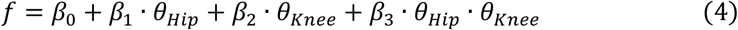

where *f* denotes the average firing frequency for one trial during the stationary period at each recording position, *θ*_*Hip*_ denotes the hip joint angle, *θ*_*Knee*_ denotes the knee joint angle, and *β*_1_~*β*_3_ denote the regression coefficients. Note that before using *θ*_*Hip*_ and *θ*_*Knee*_ as explanatory variables in the multiple regression analysis, we calculated their correlation to confirm whether multicollinearity existed between the variables. Before using *θ*_*Hip*_ and *θ*_*Knee*_ as variables in the multiple regression analysis model, they were centered by subtracting the mean value across all 16 recording positions, and then normalized to have the maximum value of 1 and the minimum value of −1. This process mitigated the effect of multicollinearity. *θ*_*Hip*_ ∙ *θ*_*Knee*_ is an interaction term that indicates the interaction between the hip and knee joint angles. The regression model used data from five trials for each of the 16 recording positions (total 80 data points).

Neurons satisfying *R*^2^ > 0.4, *p* < 0.001 in Eq. 4 were considered as neurons that responded to joint angles. The angular factors that contribute most to the activity of these neurons was evaluated using the following *r*^2^ (Bosco & Poppele, 1997).

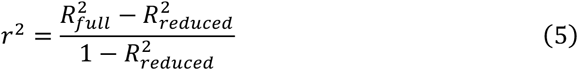

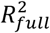 is the *R*^2^ of the regression model containing all variables (*θ*_*Hip*_, *θ*_*Knee*_, *θ*_*Hip*_ ∙ *θ*_*Knee*_) in Eq. 4, and 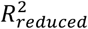 is the *R*^2^ of the regression model excluding the variable used to confirm the contribution. The statistical significance of the calculated *r*^2^ was evaluated by calculating the F-value using an incremental t-test and determining the p-value from its F-distribution. The F-value is defined by the following equation

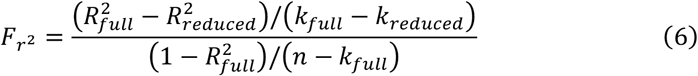

where *k*_*full*_ denotes the total number of coefficients in the regression model containing all variables in Eq. 4, *k*_*reduced*_ denotes the total number of coefficients in the regression model excluding the variables tested, and n denotes the sample size. The conditions for determining contribution of the hip joint, knee joint, and two-joint interaction were defined as follows:

Hip joint:

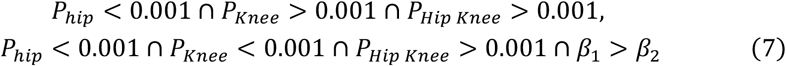

Knee joint:

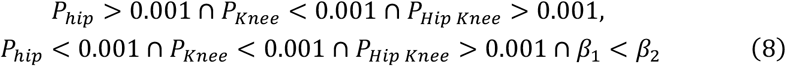

Hip and Knee joint:

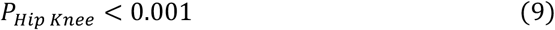

*P*_*Hip*_ denotes the p-value calculated from 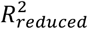 with *θ*_*Hip*_ = 0, *P*_*Knee*_ denotes the p-value calculated from 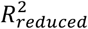 with *θ*_*Knee*_ = 0, and *P*_*Hip Knee*_ denotes the p-value calculated from 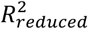 with *θ*_*Hip*_ ∙ *θ*_*Knee*_ = 0.

### Neural activity specific to a particular posture

Among the neurons that did not respond to joint angles, some exhibited activity that was specifically modulated only at a single endpoint. To identify such neurons, we conducted the following statistical analysis using the average firing rate during the stationary period of each trial (n = 5 trials per position) at each recording position as the dependent variable. Prior to this analysis, neurons that responded to joint angles were excluded.

First, a one-way analysis of variance (ANOVA) was performed with the recording position (16 levels) as the factor to test whether neural activity differed significantly across recording positions. Next, if a significant difference was observed by the ANOVA, a post-hoc Tukey-Kramer test was conducted to identify recording positions with significantly different activity compared to the others. For each recording position, we counted the number of significant pairs (p < 0.05) between that position and the other 15 other recording positions, and then calculated the mean number and standard deviation (SD) of these counts across all 16 recording positions. Finally, a neuron was classified as showing selective modulation at a single endpoint if, at one specific recording position, the number of significant pairs exceeded the mean +3SD and was 8 or more (i.e., more than half of the total 15 comparison pairs).

### Neural Population Analysis by Principal Component Analysis

To examine how populations of spinal neurons responded to change in hindlimb posture, we performed principal component analysis (PCA). Prior to PCA, neural activity was normalized as follow (Sharma *et al*., 2024). For each trial, the firing rate was calculated every 50 ms during the stationary period at each recording position. Then, for each neuron, Z-score normalization was applied to the 50-ms binned firing rate across all 16 recording positions within each trial, thereby equalizing differences in activity levels among neurons. Subsequently, the normalized data were time-averaged over the stationary period for each recording position. The resulting time-averaged values were then averaged across all five trials to construct the final dataset for PCA. PCA was performed using the normalized and time-averaged neural activity as variables (columns) and the 16 recording positions as observations (rows).

Components whose cumulative contribution rate first exceeded 80% were regarded as the major activity patterns, and their characteristics were investigated. To interpret each principal component (PC), we examined how strongly and in which direction each neuron contributed to that component (i.e. loading), and related those loadings to the response characteristics of individual neuron identified in the analyses described above, i.e., “Joint angle encoding by single neurons” and “Specific endpoint encoding by single neurons”. The loadings represent both the magnitude and direction of each neuron’s projection onto the basis vector in the PC space.

To quantitatively analyze those loadings, we first normalized the value for each neuron so that the sum of the squared loadings for each PC equaled 1. Each neuron was then categorized into five response types: those responsive to the hip joint, knee joint, both hip and knee joints, those specifically responsive to a single endpoint, and those unresponsive to postural changes. The normalized loadings were then summed within each response category. This procedure allowed for a quantitative assessment of the proportion of each response type contributing to each principal component.

### Endpoint Decoding from Measured Neural Activity

To examine whether the activity of recorded spinal neurons contained sufficient information to represent posture, we performed a decoding analysis to predict posture from the neural activity. Neural activity was normalized for each trial using the same method as that described in the *Neural Population Analysis by Principal Component Analysis*. The postural information to be decoded was defined as the orientation (O) of the leg axis (the straight line connecting the hip and ankle joint) relative to the vertical, and the length (L) of the leg axis (Lacquaniti & Maioli, 1994). Data from 5 trials recorded at each of the 16 recording positions (a total of 80 data points) were used for decoding. Lasso regression (Tibshirani, 1996), a form of sparse linear regression, was employed for the decoding analysis.

To evaluate decoding performance, which reflects the degree to which neural activity represents posture, cross-validation was performed using data from 15 recording positions as training data and the remaining one as test data. The regression model was optimized by minimizing the following objective function:

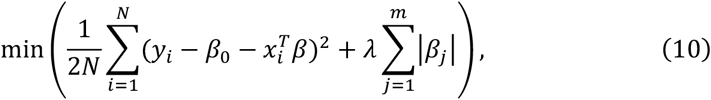

where *N* denotes the number of training data (N=75), *m* denotes the total number of recorded neural activities, *y*_*i*_ denotes the endpoint parameters (O and L) for the *i*-th training sample, and *x*_*i*_ denotes *m* neural activities for the *i*-th training data. *β*_0_ is the intercept, *β* is the regression coefficient, and *λ* is the regularization parameter. The regularization parameter *λ* was determined through 5-fold cross-validation within the training data to minimize the objective function.

After training model using 16 combinations of data (15 positions for training and 1 position for testing), the predicted endpoint parameters for each test recording position were computed. As a final measure of decoding performance, the coefficient of determination (*R*^2^) was calculated by integrating all prediction results from five trials recorded at each of the 16 recording positions.

## Results

Recordings were obtained from a total of 208 spinal neurons along a single electrode track in two cats (Cat A and Cat B), comprising 72 neurons in Cat A and 136 neurons in Cat B. The results of subsequent analyses are presented in the following order: (1) analyses of single-neuron activity and (2) analyses of simultaneously recorded neuronal populations.

### Different static hindlimb postures modulate spinal neuronal activity

We first imposed 16 preprogrammed static recording positions of the hindlimb using a robotic manipulator. For each position, we verified the two-dimensional (X–Y) locations of the iliac landmark, hip, knee, and ankle by video-based motion analysis (Figure 2B). From these measurements, the limb configurations corresponding to the 16 recording positions were estimated (See Figure 2D for Cat A, and S1 for Cat B).

Using the estimated landmarks, we computed hip and knee joint angles for each recorded posture according to the definitions in Figure 2C. To assess potential multicollinearity in subsequent multiple-regression analyses, we quantified the linear correlation between hip and knee angles across the sampled postures. The correlation was not significant (r = 0.18, p = 0.51), indicating minimal collinearity between regressors and supporting the reliability of the model.

We next compared firing rates of spinal neurons across the 16 recording positions. Figure 4 shows a representative neuron whose activity covaried with Y-axis (proximal–distal) coordinate of the hindlimb position: its firing rate increased at more proximal positions and decreased at more distal positions.

**Figure 4.**
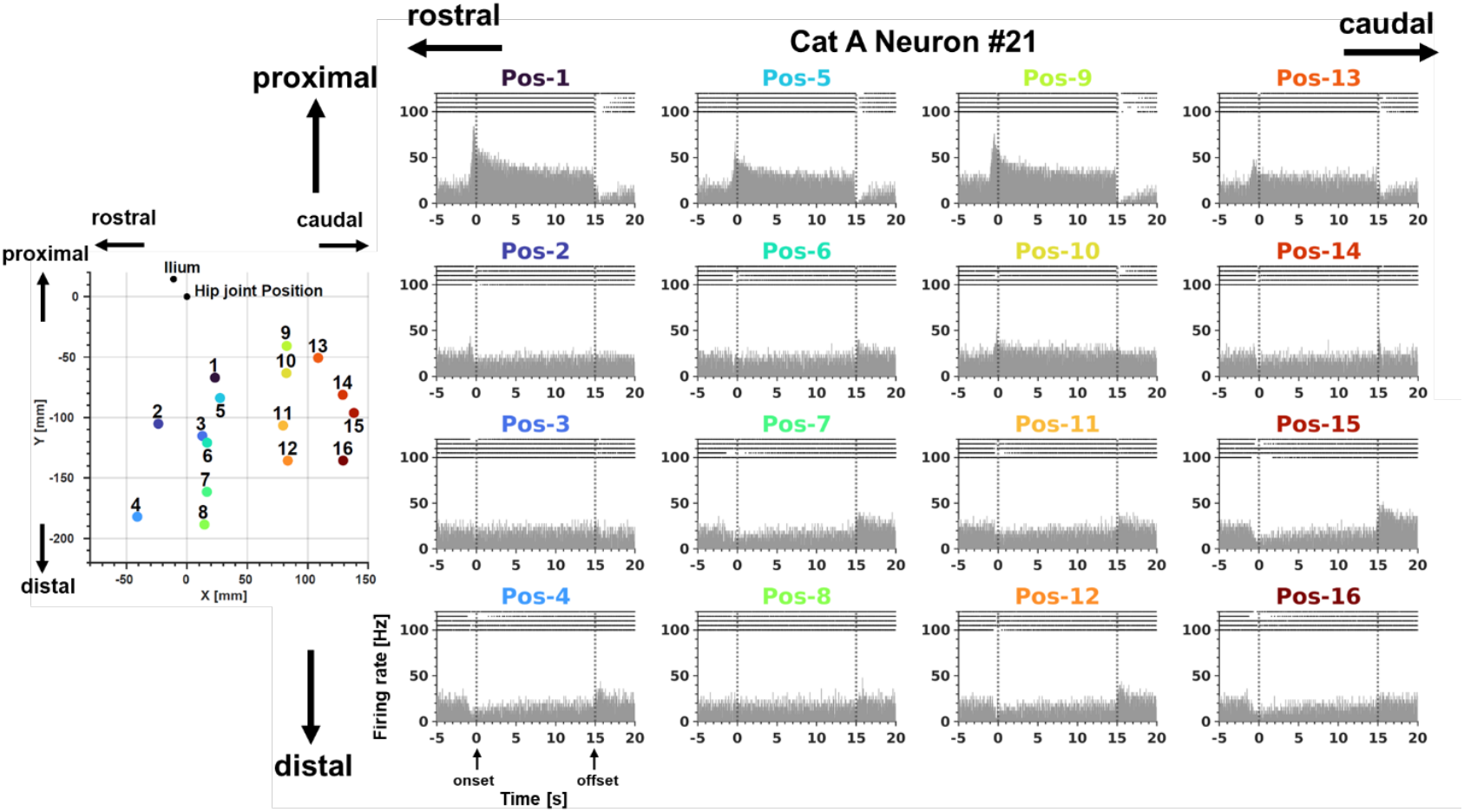
Neural activity (Cat A Neuron #21) modulated by hindlimb position. The right panels illustrate the firing frequency for each corresponding recording position, and the details of the figure are as described in Figure 3A. Each panel of neural activity is arranged according to the cat’s ankle position at the recording location: proximal positions are at the top, distal positions are at the bottom, rostral positions are on the left, and caudal positions are on the right. The cat’s ankle location for each recording position is indicated by a colored dot in the left panel. The same colors are used to label the corresponding recording positions in the right panels.

Population summaries for Cat A are shown in Figure 5, where each panel is arranged on a grid according to the corresponding X–Y location of the recording position. Consistent with the exemplar in Figure 4, several neurons exhibited posture-dependent modulation across the recorded configurations. For example, neuron #53 (▲) varied primarily with the endpoint’s X-axis coordinate, whereas neuron #41 (■) covaried with the Y-axis coordinate, similar to neuron #21 (the neuron shown in Figure 4). Comparable patterns were observed in Cat B (Figure S2).

**Figure 5.**
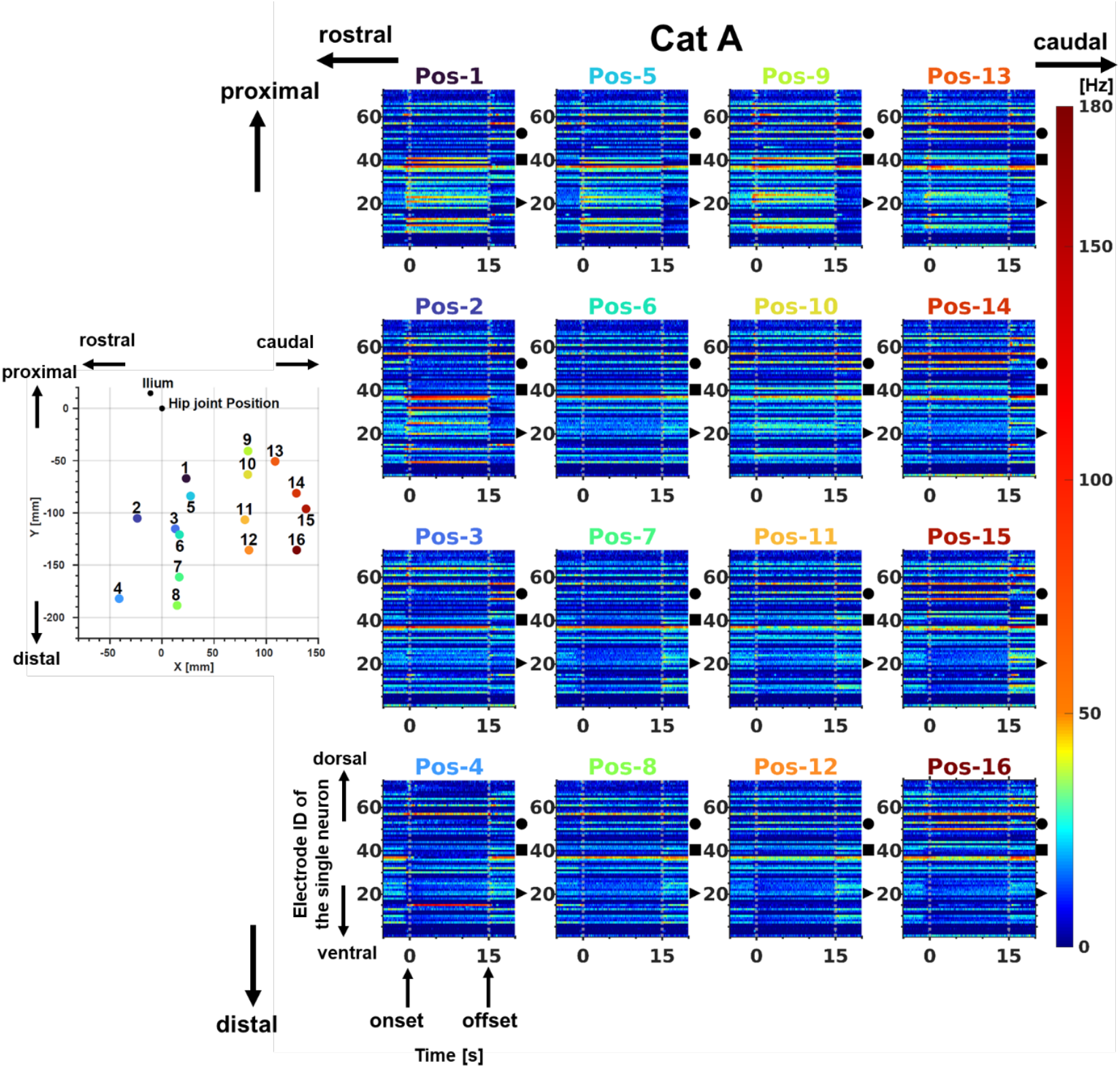
Activity of 72 single neurons across 16 recording positions in Cat. **A**. The right panels show the neural activity of 72 single neurons for each corresponding recording position; other details are as described in Figure 3B. The spatial arrangement of the right panels follows the configuration shown in Figure 4. Markers on the vertical axis denote specific neurons: ▸Neuron #21(shown in Figure 3), ■ Neuron #41 (responding to changes in the endpoint’s Y-axis), and ● Neuron #53 (responding to changes in the endpoint’s X-axis).

### Relationship between single-neuron firing rate modulation and joint-angle variables

To test whether the firing-rate modulation observed in Figures 5 and S2 reflected joint-angle information, we performed multiple linear regression with hip and knee joint angles—both systematically varied by the recording positions—as predictors. In Cat A, 23/72 neurons (31.9%) showed significant covariation with joint angles (R^2^ > 0.40, p < 0.001). In Cat B, 12/136 neurons (8.8%) met the same criterion (R^2^ > 0.40, p < 0.001). Representative neurons and fitted regression functions are shown in Figure 6. In Cat A, neuron #53 (Figure 6A) and in Cat B, neuron #64 (Figure 6B) increased firing as the hip became more flexed. For knee-tuned neurons, Cat A neuron #21 (Figure 6C) fired more at flexed knee positions, whereas Cat B neuron #81 (Figure 6D) increased firing toward knee extension. A neuron responsive to both joints in Cat A (neuron #24; Figure 6E) exhibited maximal activity for a coordinated combination of hip extension with knee flexion.

**Figure 6.**
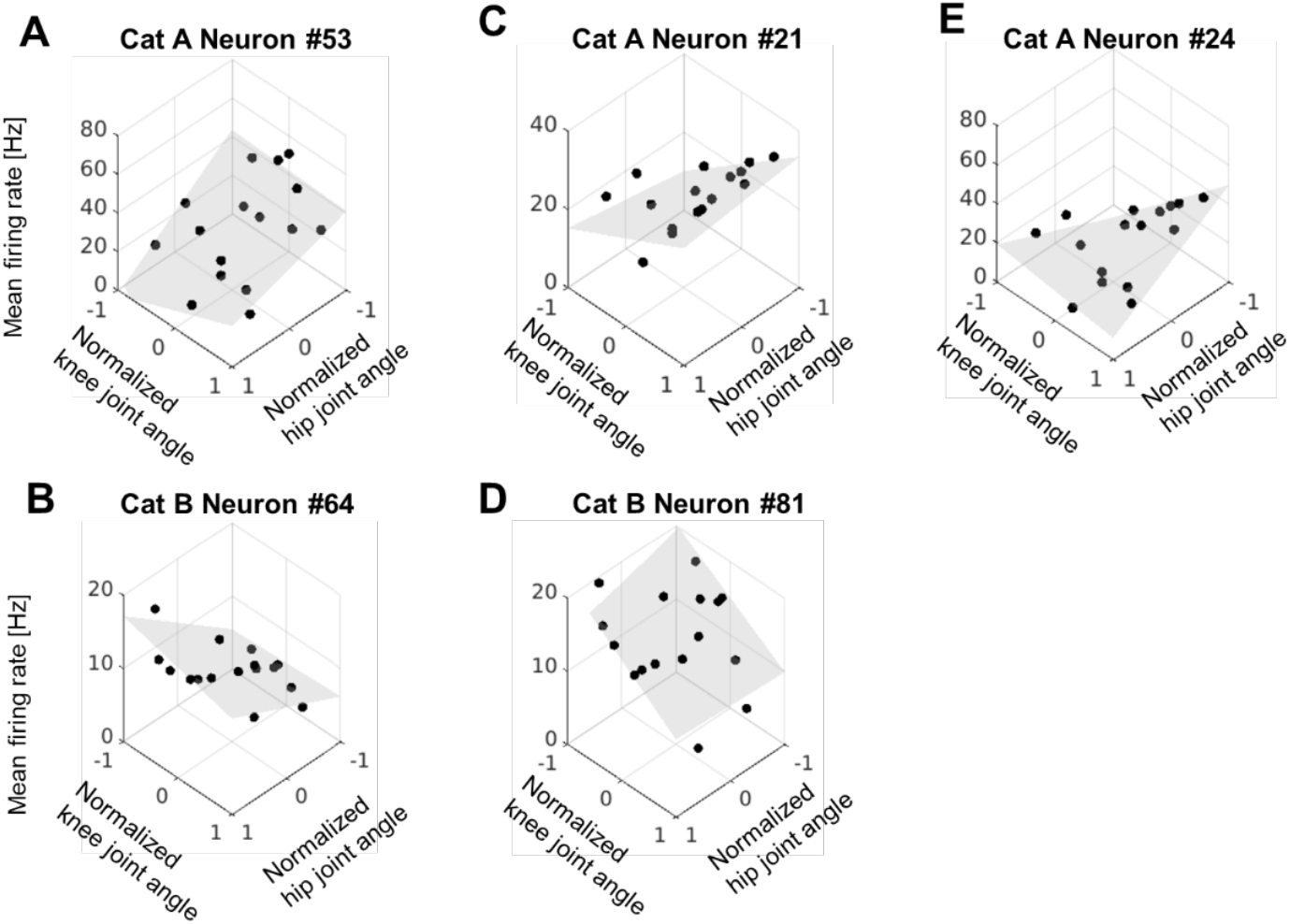
Neurons modulated by joint angles. The X-axis represents the normalized and centered hip joint angle, the Y-axis represents the normalized and centered knee joint angle, and the Z-axis represents the average firing frequency. The dots in the figure represent the average firing frequency during the stationary period across 5 trials. The gray surface is the predicted average firing frequency estimated by the multiple regression model. **A**, Neuron #53 responsive to the hip joint in Cat A: *f* = 22.87 + 4.20 ∗ *θ*_*Hip*_ − 0.19 ∗ *θ*_*Knee*_ − 2.17 ∗ *θ*_*Hip*_ ∗ *θ*_*Knee*_; *R*^2^ = 0.403.**B**, Neuron #64 responsive to the hip joint in Cat B: *f* = 10.43 + 4.65 ∗ *θ*_*Hip*_ − 0.84 ∗ *θ*_*Knee*_ − 1.20 ∗ *θ*_*Hip*_ ∗ *θ*_*Knee*_; *R*^2^ = 0.605.**C**, Neuron #21 responsive to the knee joint in Cat A: *f* = 22.07 + 0.40 ∗ *θ*_*Hip*_ + 9.53 ∗ *θ*_*Knee*_ − 2.20 ∗ *θ*_*Hip*_ ∗ *θ*_*Knee*_; *R*^2^ = 0.584. **D**, Neuron #81 responsive to the knee joint in Cat B: *f* = 15.78 + 0.73 ∗ *θ*_*Hip*_ − 5.57 ∗ *θ*_*Knee*_ − 0.38 ∗ *θ*_*Hip*_ ∗ *θ*_*Knee*_; *R*^2^ = 0.666. **E**, Neuron #24 responsive to the hip and knee joint in Cat A: *f* = 18.39 − 3.31 ∗ *θ*_*Hip*_ + 11.89 ∗ *θ*_*Knee*_ − 16.08 ∗ *θ*_*Hip*_ ∗ *θ*_*Knee*_; *R*^2^ = 0.533.

Among the joint-angle–responsive neurons in Cat A, 6/23 (26.1%) were selectively related to hip angle, 10/23 (43.5%) to knee angle, and 7/23 (30.4%) to both angles. In Cat B, 10/12 (83.3%) covaried with hip angle and 2/12 (16.7%) with knee angle; no neurons in Cat B exhibited joint tuning to both angles.

### Spinal neurons selectively responsive to a single leg configuration

We next analyzed neurons that did not show covariation with joint angles in the regression analysis (Cat A: 49/72; Cat B: 124/136). We frequently observed that, despite lacking systematic dependence on hip or knee angles, some neurons exhibited marked modulation at a single hindlimb posture (i.e., one endpoint position) only. An example is shown in Figure 7: Cat A, neuron #15 showed no linear covariation with the endpoint’s X- or Y-axis coordinates, yet displayed a pronounced increase in firing at Pos-4. Specifically, its mean firing rate at Pos-4 across five trials was 103.5 Hz, which was substantially higher than the range observed at the other positions (0.9–22.1 Hz).

**Figure 7.**
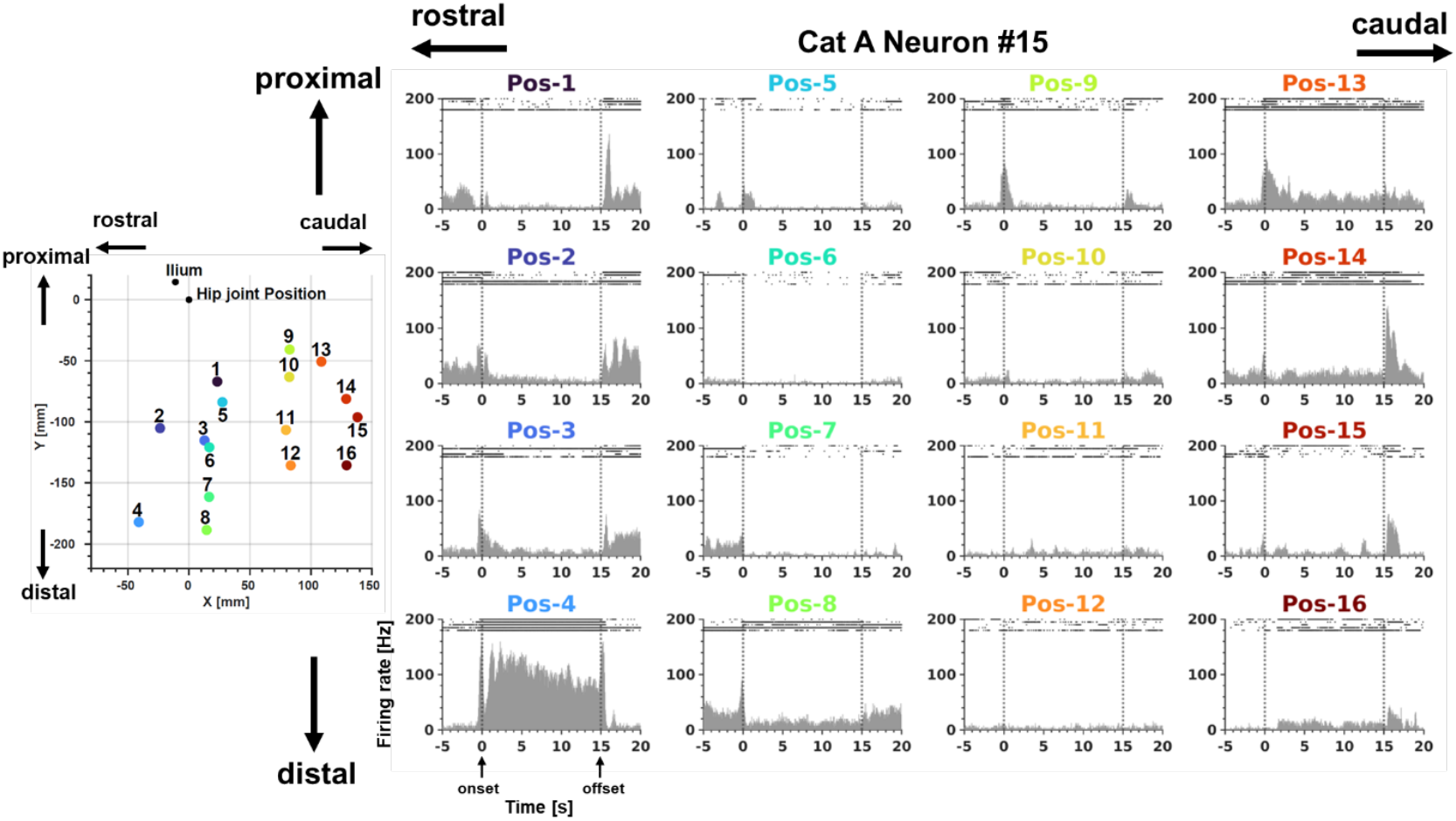
Neural activity (Cat A, Neuron #15) showing a large increase at a single recording position. The right panels show the firing frequency for each corresponding recording position; other details are as described in Figure 3A. The spatial arrangement of the right panels is identical to that shown in Figure 4.

We therefore tested all non–joint-angle–responsive neurons for posture selectivity based on multiple-comparisons procedures (see *Neural activity specific to a particular posture* in Methods). Significant selectivity for a single position was observed in 11/49 neurons (22.4%) in Cat A and 7/126 neurons (5.5%) in Cat B. In Cat A, the selected positions were distributed as follows: Pos-1, 3/11 (27.3%); Pos-2, 5/11 (45.5%); Pos-4, 2/11 (18.1%); and Pos-14, 1/11 (9.1%). In Cat B, selective neurons were observed at Pos-4, 3/7 (42.9%); Pos-9, 2/7 (28.6%); and Pos-16, 2/7 (28.6%).

Notably, a majority of these posture-selective neurons were maximally active at configurations in which hip and/or knee angles reached extreme flexion or extension within the sampled workspace (Cat A: 11/11; Cat B: 5/7).

### Population representation of joint angles and leg configuration

Up to this point, we have analyzed firing rate modulations at the single-neuron level, as in previous studies (Bosco *et al*., 1996). A key distinction of the present study is that we recorded from many spinal neurons simultaneously while the hindlimb was held at systematically varied static postures, enabling population analyses that are not possible with serial, single-unit-based approaches. Using z-scored mean firing rates across the 16 endpoint positions, we therefore performed principal component analysis (PCA).

In Cat A, PC1, PC2, and PC3 explained 46.4%, 20.6%, and 13.6% of the variance, respectively, accounting for a cumulative variance of 80.8% (Figure 8A). The distribution of PC scores across the 16 postures (Figure 8B) revealed a functionally organized structure: PC1 scores were higher for postures with a flexed knee and lower for those with extended knee; PC2 scores were higher for hip flexion and lower for hip extension; and PC3 showed a sharp positive excursion at Pos-4 and a marked negative excursion at Pos-2.

**Figure 8.**
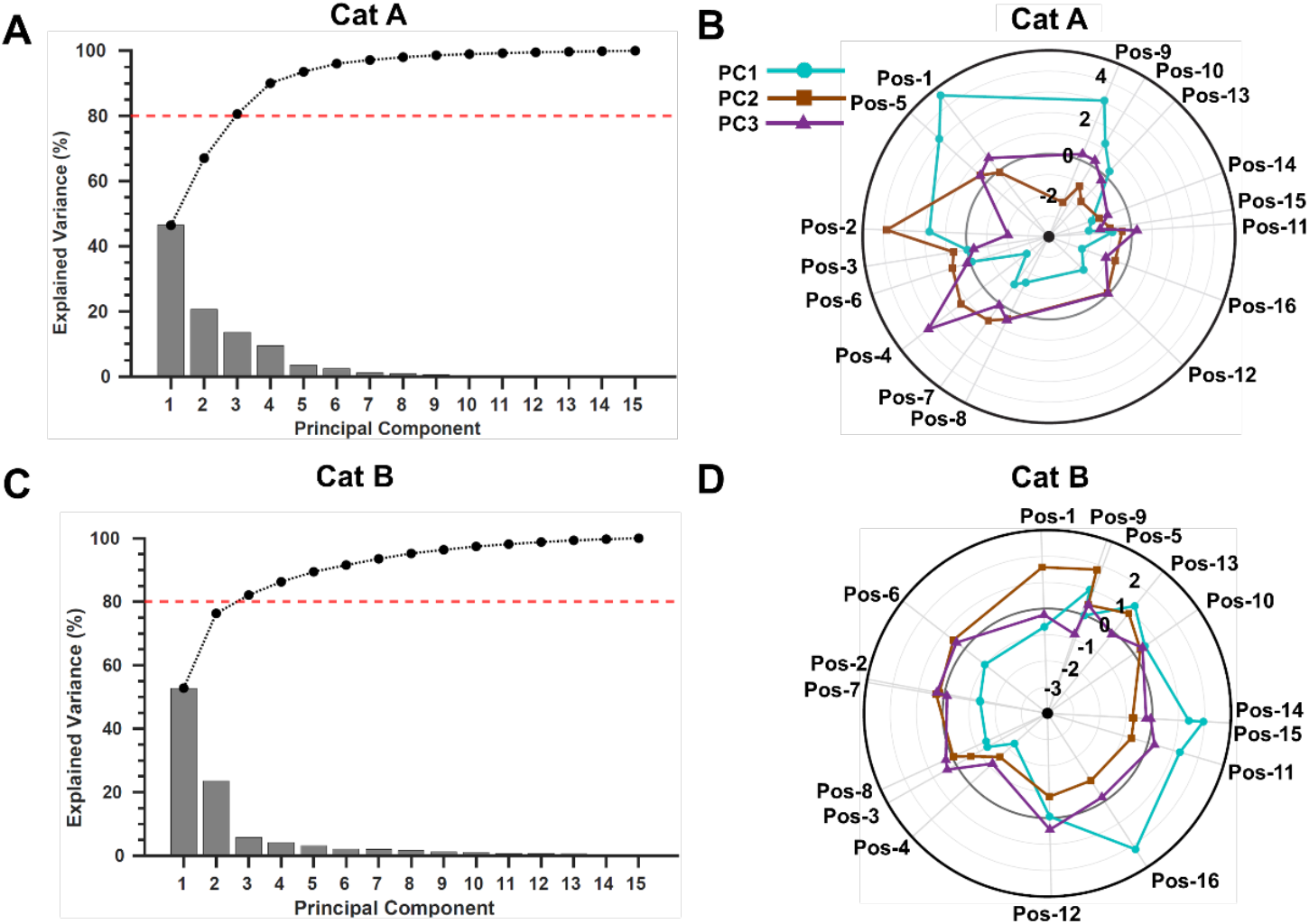
Contribution rates and PC scores calculated by PCA. **A**, Principal component contribution rates for Cat A. The bar graph shows the contribution rate of each component, and the line graph at the top represents the cumulative contribution rate. **B**, PC scores for each recording position in Cat A. Each recording posture was arranged in polar coordinates based on the angle information obtained after transformation into polar coordinates, centered on the mean X–Y coordinates of the ankle joint. The radial axis represents the PC score. PC1 is shown by blue circles (●), PC2 by brown squares (■), and PC3 by purple triangles (▲). **C**, Principal component contribution rates for Cat B. Data representation is the same as in A. **D**, PC scores for each recording position in Cat B. The data representation is the same as in B.

In Cat B, PC1, PC2, and PC3 explained 52.8%, 23.5%, and 5.8% of the variance (cumulative 82.1%; Figure 8C), accounting for most of the total variance. PC scores (Figure 8D) again aligned with joint configuration: PC1 increased with hip extension and decreased with hip flexion; PC2 increased with knee flexion and decreased with knee extension; PC3 exhibited comparatively low scores at Pos-4 and Pos-9 relative to the other postures.

To relate these principal axes to the single-neuron tuning classes, we associated each neuron’s normalized loadings on PC1–PC3 with its independently defined response category (hip-tuned, knee-tuned, bi-joint-tuned, single-endpoint-selective, or nonresponsive; see *Neural Population Analysis by Principal Component Analysis* in Methods). We then computed, for each PC, the proportion contributed by neurons from each category.

In Cat A (Figure 9A), knee-tuned neurons predominated in PC1 (59.5%). For PC2, hip-tuned and bi-joint neurons contributed 30.7% and 21.5%, respectively. In PC3, single-endpoint-selective neurons accounted for 53.1%. In Cat B (Figure 9B), hip-tuned neurons dominated PC1 (53.1%), and knee-tuned neurons comprised ~half of PC2 (48.5%). For PC3, nonresponsive neurons formed the largest share (68.5%); excluding these, single-endpoint selective neurons contributed 17.9%, exceeding any individual joint-tuned category.

**Figure 9.**
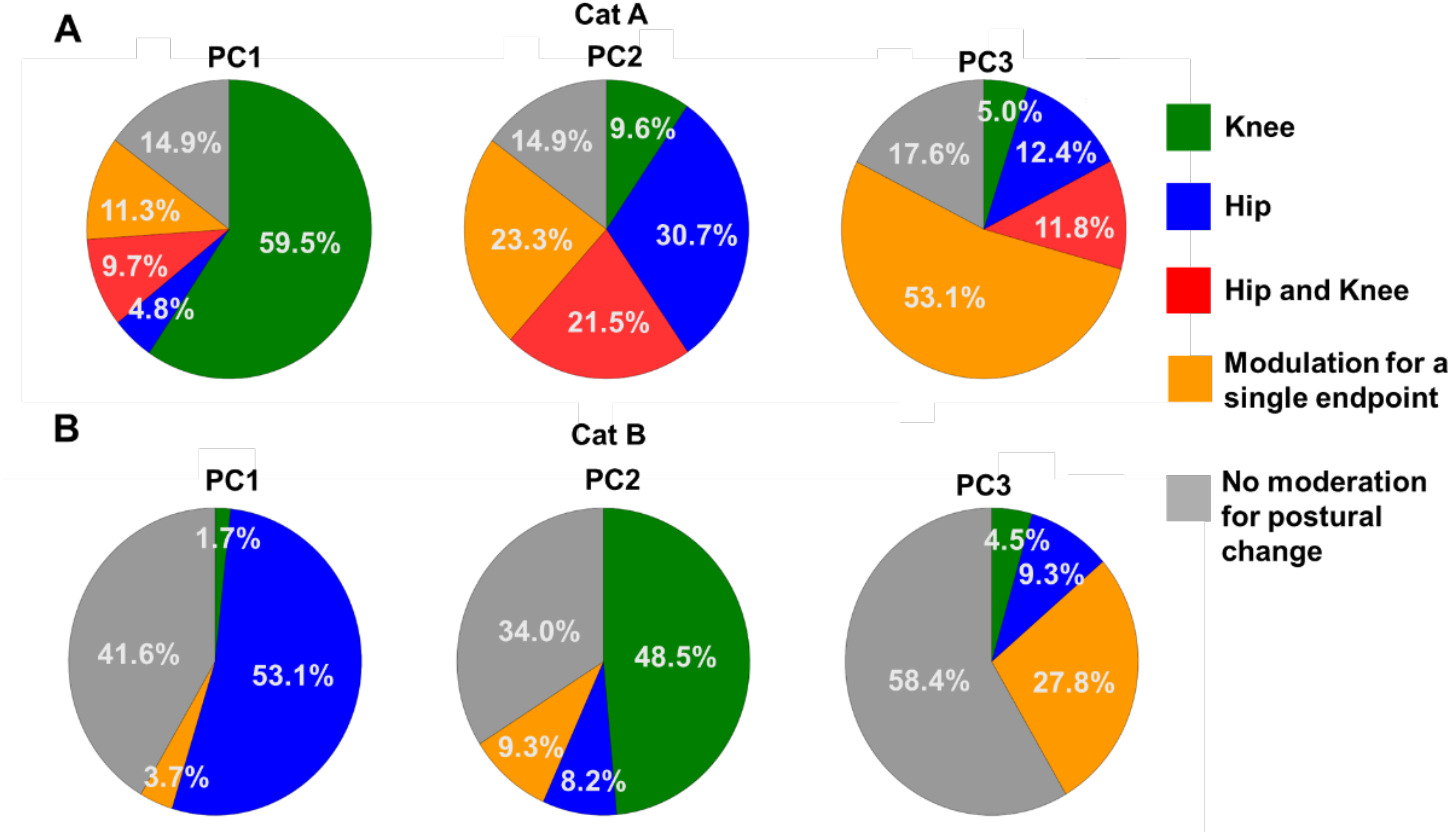
Distribution of response characteristics related to joint angles and specific endpoint encoding on the neuronal loadings that constitute each principal component (PC1–PC3). A, Distribution of response characteristics for each PC in Cat A. Green indicates neurons responsive to the knee joint, blue indicates neurons responsive to the hip joint, red indicates neurons responsive to both the knee and hip joints, orange indicates neurons responsive to a specific endpoint, and gray indicates neurons nonresponsive to postural changes. B, Distribution of response characteristics for each PC in Cat B. The data presentation is the same as in A.

Collectively, the leading population axes captured (i) single-joint angle variation (knee for Cat A PC1; hip for Cat B’s PC1), (ii) the complementary joint (hip and bi-joint for Cat A’s PC2; knee for Cat B’s PC2), and (iii) posture-specific patterns not explained by continuous joint-angle changes (PC3). These results indicate that the spinal ensemble in this work comprised neurons representing hip or knee angles individually (Cat A and Cat B), their joint combinations (Cat A), and specific endpoint configurations (Cat A and Cat B).

#### Endpoint decoding by spinal neurons

Next, decoding analysis was performed using LASSO regression (Sparse linear regression) to investigate whether the endpoint could be predicted using the recorded neural population activity. The polar coordinate system (Orientation and Length), based on the hip joint position, was adopted as the metric representing the endpoint and was defined as the parameter to be predicted in the decoding analysis (see *Endpoint Decoding from Measured Neural Activity* in Methods).

The predicted endpoint values estimated by the model using 72 neurons and 15 of the 16 recording positions (with one position excluded) in Cat A are shown in Figure 10A for Orientation and Figure 10B for Length. The *R*^2^ values indicating the prediction accuracy of the model were *R*^2^ = 0.931 for Orientation and *R*^2^ = 0.782 for Length. These values were high compared to previous decoding studies (Fathi & Erfanian, 2022).

**Figure 10.**
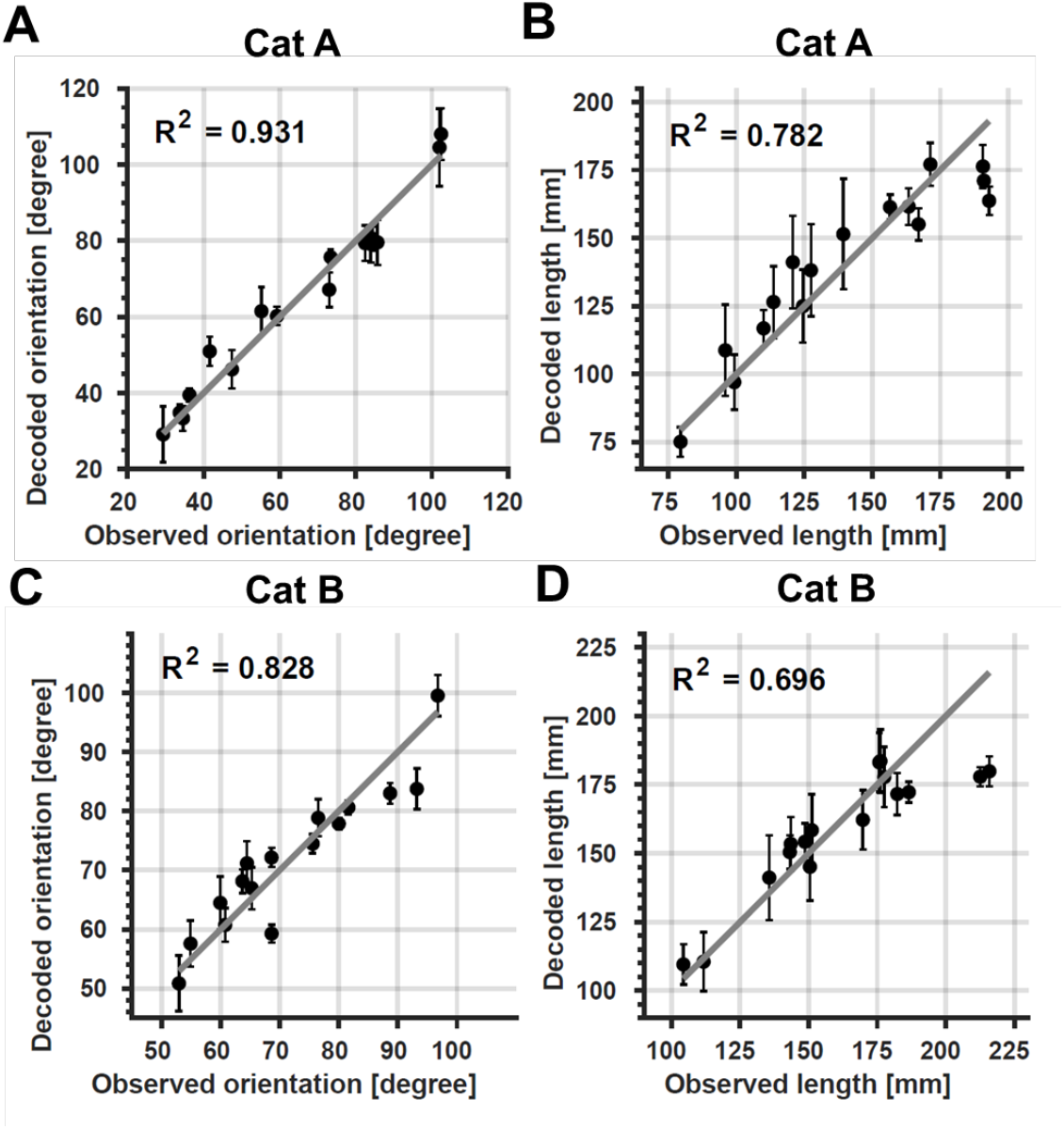
Comparison of predicted and observed endpoint values from neural population decoding. **A**, Predicted orientation values for Cat A. The horizontal axis represents the observed value, and the vertical axis represents the predicted value from decoding. Black circles (●) show the mean of the five predicted values at each recording position, and error bars indicate the standard deviation (SD). The gray line represents the ideal prediction line (observed value = predicted value; slope of 1). **B**, Predicted length values for Cat A. **C**, Predicted orientation values for Cat B. **D**, Predicted length values for Cat B. Axis labels, data representation, and symbols used in panels B-D are the same as in A.

The predicted values from the model created using 136 neurons in Cat B are shown in Figure 10C for Orientation and Figure 10D for Length. The R^2^ values indicating the prediction accuracy of the model were *R*^2^ = 0.830 for Orientation and *R*^2^ = 0.694 for Length, which also demonstrated favorable prediction performance.

## Discussion

Here, we simultaneously recorded lumbosacral spinal neurons in decerebrate, immobilized cats while a robot maintained the hindlimb at 16 static endpoint positions spanning hip–knee configurations. A subset of neurons covaried significantly with joint angles, with tuning distributed across hip-, knee-, and bi-joint sensitivities, whereas another subset was selectively active at a single endpoint configuration despite lacking continuous joint-angle dependence. Population analyses (PCA) revealed physiologically meaningful structure: in both animals, PC1 and PC2 tracked knee and hip angle, respectively, while PC3 captured a single end-point-specific pattern not explained by continuous angle changes. Decoding analyses further indicated that whole-limb kinematic parameters (Bosco & Poppele, 1993), such as hindlimb endpoint position, can be reconstructed by integrating signals from a relatively small subset of spinal neurons. Together, these results show that spinal ensembles encode static limb configuration through a mixture of single-joint, joint-combination, and configuration-specific representations.

### Possible hierarchical organization of proprioceptive representation in the spinal cord

It is well established that proprioceptors and primary afferents encode muscle length and single joint angles (Grigg, 1975; Grigg & Greenspan, 1977; Ferrell, 1980; Wei *et al*., 1986; Proske & Gandevia, 2012). In contrast, studies of the dorsal spinocerebellar tract (DSCT)—the major ascending pathway originating from the spinal cord—have demonstrated that DSCT neurons encode limb posture through multiple mechanisms (Bosco *et al*., 1996; Bosco *et al*., 2000). When the hindlimb is passively positioned, approximately 67% of DSCT neurons are modulated by two or more joint angles, indicating convergence of sensory information from multiple joints rather than single-joint representations (Bosco *et al*., 1996). In other words, proprioceptive signals from diverse peripheral receptors converge onto DSCT neurons, enabling the encoding of endpoint position in limb-centered coordinates, which is then transmitted to higher centers. In this study, recording has been made from “non-DSCT” neurons localized the outside of spinal segment for the DSCT (T9-L4 segments) (Rexed, 1952; Tanaka & Hirai, 1994; Veshchitskii *et al*., 2022). The non-DSCT spinal neurons in the present study appear to represent an intermediate stage between the primary afferents and DSCT neurons. Among the non-DSCT population, most neurons represented the variance of a single joint angle. In Cat A, the first principal component (PC1) covaried with knee angle, and 59.5% of its loadings were accounted for by the “knee neurons” identified in the single-unit analysis. In Cat B, the first principal component covaried with hip angle, with 53.1% of its loadings explained by the “hip neurons.” Thus, the majority of spinal neurons, like primary afferents, encode single-joint information, whereas a smaller subset exhibited multi-joint modulation similar to that observed in DSCT neurons.

These findings may suggest a hierarchical organization of proprioceptive representation within the spinal cord: transformations from a joint-centered to a limb-centered coordinate frame may occur progressively, from receptor-level afferents, through interneuronal populations, to ascending projection neurons such as DSCT cells. Supporting this view, studies of the rat external cuneate nucleus have shown that neuronal activity correlates strongly with limb-axis length and orientation, but only weakly with individual joint angles (Giaquinta et al., 1999). Together, these results suggest that proprioceptive information is integrated hierarchically within the spinal cord before being relayed to higher-order sensorimotor structures.

However, in addition to the DSCT, there are ascending pathways that originate from lumbar spinal interneurons—most notably postsynaptic dorsal column (PSDC) neurons (Rustioni & Kaufman, 1977; Brown & Fyffe, 1981) —which also convey hindlimb proprioceptive information to the medulla and ultimately to the cerebral cortex. Given that cortical proprioceptive representations are organized across combinations of multiple joints (Klaes *et al*., 2015; Goodman *et al*., 2019; Chowdhury *et al*., 2020), it is possible that a subset of the cells we recorded whose activity covaried with changes in two joint angles were in fact PSDC neurons. The proprioceptive representations of spinal neurons in ascending pathways other than the DSCT remain largely unexplored; future studies are needed to address this.

### Neurons representing single endpoints

In both animals, we identified a third principal component composed predominantly of neurons that were selective for a single endpoint configuration—a specific combination of hip and knee angles. These preferred configurations were located near the limits of the sampled joint workspace. To our knowledge, such single endpoint tuning at the spinal level has not been reported previously, suggesting a potentially novel type of proprioceptive representation.

DSCT neurons are known to encode hindlimb posture in a limb-centered coordinate system, rather than a joint-based one (Bosco *et al*., 2000). In contrast, the single-endpoint neurons described here exhibited a discrete, posture-specific selectivity: their firing rates changed significantly only at one specific endpoint, without a graded dependence across neighboring postures. Thus, unlike DSCT neurons that covary continuously with endpoint position along the limb-centered coordinate, these neurons encode categorical information confined to a single posture.

Some joint-receptor afferents show similar response patterns, being relatively flat across midrange joint angles and increasing activity only near movement limits (Clark & Burgess, 1975; Grigg, 1975). However, these afferents respond near the extreme positions of a single joint, whereas the neurons identified in the present study (e.g. Figure 7) were tuned to a specific combination of two joint angles—a feature not previously described in spinal or supraspinal proprioceptive neurons.

One possible reason why such neurons were not reported in earlier studies may relate to differences in workspace coverage. The present study systematically varied both the X and Y components of the endpoint position, covering approximately 45.6 % of the hip and 58.8 % of the knee range of motion for Cat A, and 18.4 % of the hip and 56.3 % of the knee for Cat B (Jaeger *et al*., 2007). In contrast, previous DSCT experiments explored a relatively narrow range of hindlimb positions (as illustrated in Bosco *et al*., 1996, Figure 1). This broader sampling may have been essential for detecting posture-specific peaks in neural activity that would be missed in narrower paradigms. Another possibility is that such selective neurons do not project through the DSCT pathway—that is, this category of non-DSCT neurons may represent local spinal processing that is not relayed to supraspinal centers via the DSCT but via other ascending pathways (see discussion). Direct testing of this possibility will require experiments recording from DSCT and other ascending spinal neurons under a comparable posture-sampling paradigm.

The potential functional significance of single-endpoint neurons remains speculative, but several interpretations are plausible. First, these interneurons may act as landmark detectors for specific endpoint, providing discrete cues that switch motor control modes. Many behaviors are cyclic or sequential: in gait, transitions between extension and flexion phases occur near kinematic extrema (Buford & Smith, 1990; Hiebert *et al*., 1996); in reach-to-grasp task, arm posture at the end of reaching is carefully planned in advance and plays a critical role in guiding grasp initiation through both feedforward and feedback mechanisms (Grea *et al*., 2000; Sangole & Levin, 2008). Encoding these extrema at the spinal level could trigger phase changes in CPGs (Yakovenko *et al*., 2005) and facilitate smooth transit from reaching to grasping (Rouse & Schieber, 2016).

Second, these neurons may serve as boundary monitors, signaling when the limb approaches workspace extremes rather than a single-joint limit, thereby facilitating protective strategies such as co-contraction or joint stiffening. In the design of engineered systems such as robots, mechanisms for monitoring and preventing movement outside the workspace extremes are commonly employed (Standardization., 2025). In the other aspect, the boundaries of the workspace, for example in the case of a limb, represents the state in which the limb is maximally extended and capable of reaching the farthest distance, making movements at the boundary functionally effective. Thus, in control design, there is a demand for motion control near this boundary (Maciejowski, 2002). In the design of engineered systems such as robots, a common approach to fulfilling this control demand is to generate optimal control input under constraints that prevents the system from reaching the boundary (Maciejowski, 2002). Therefore, information about the boundary contributes to both mechanical safety and the efficiency of control design. As a neuronal foundation to support these, spinal single-endpoint signals could provide a compact, categorical marker of limb-centered boundary states, enabling rapid gain adjustments that complement the continuous encoding of joint angles.

Third, internal calibration landmarks: fixed “nodes” at reproducible endpoint locations could serve as reference anchors for recalibrating population codes that drift or adapt over time, thereby supporting robust proprioceptive estimates across days or changing load conditions. This idea aligns with evidence that proprioceptive maps recalibrate after cross-sensory discrepancies and that such changes persist over hours to days (Ostry *et al*., 2010; Tsay *et al*., 2022). It also echoes findings that, despite day-to-day population drift, stable readout can be maintained by aligning neural activity to consistent low-dimensional structure across sessions (Degenhart *et al*., 2020; Rule *et al*., 2020). Furthermore, during motor learning under novel dynamics the CNS can relearn and recalibrate internal models when task mechanics change (Shadmehr & Mussa-Ivaldi, 1994), consistent with the notion that endpoint “anchors” could facilitate rapid re-centering of spinal population codes under varying loads.

Taken together, the discovery of single-endpoint neurons suggests that the spinal cord incorporates both continuous and categorical coding schemes: continuous representations for the majority of the workspace, and categorical boundary signals that enhance stability and safety of movement control near the limits of limb motion.

### End-point position can be decoded from the activity of spinal interneuronal populations

Analysis of firing modulation across different hindlimb endpoint positions, together with principal component analysis, revealed that the recorded spinal neurons primarily encoded proprioceptive information in a joint-based coordinate frame. As discussed earlier, this differs from the dominant coding schemes observed in DSCT neurons and cortical neurons, where proprioceptive representations are largely expressed in body- or limb-centered coordinates rather than single-joint space. Many DSCT neurons receive their inputs polysynaptically through spinal interneurons (the so-called polysynaptic pathway (Bosco & Poppele, 2001)). Therefore, joint-based representations observed in non-DSCT spinal interneurons may be integrated and transformed by ascending projection neurons—such as DSCT or PSDC cells—into a higher-dimensional, limb-centered representation.

Our decoding analyses demonstrated that the collective activity of L6 spinal neurons (n = 72 or 136) contained sufficient information to predict the endpoint position of the limb in body-centered coordinates. This finding indicates that, even at the spinal level, ensembles of non-DSCT interneurons encode sufficient information to reconstruct whole-limb kinematics. Consequently, higher-order sensory relay nuclei such as the DSCT and gracile nucleus may be capable of performing coordinate-frame transformations based on inputs from a relatively small subset of spinal interneurons via polysynaptic pathways.

Interestingly, the prediction accuracy for the length in one animal (Cat B) was poor specifically at the end-point position used for our recording with the greatest (Pos-4) and the second greatest length (Pos-16) compared to other positions (Figure 10D). This result may reflect the fact that the sparce regression model, including Lasso regression model used in this study, might fail to capture the nonlinear activity profile of single-endpoint neurons. For example, among seven single-endpoint neurons in this animal, the corresponding end-point position of five neurons (5/7, 71.4%) were either Pos-4 (three) or Pos-16 (two). In the sparce regression model, the coefficient for the variables that less contributing to the modeling are assigned as zero (Tibshirani, 1996), and, in our analysis, the coefficient of six out of seven (6/7, 85.7%) were indeed assigned as zero. This result strongly suggest that the single-endpoint neurons did not contribute to the model. Therefore, the model primarily relied on neurons showing a continuous response to endpoint changes, and essentially excluded the neurons showing categorical responding to a single endpoint. This result also suggest that the spinal cord may use two distinct coding schemes for limb proprioception; the linear, continuous coding for the endpoint changes, and nonlinear, categorical coding for the specific end position.

### Limitations

Our study has several limitations. First, in each cat we obtained recordings from only a limited number of penetrational tracks in the lumbar cord (Cat A: L5, L6; Cat B: L6). However, only the L6 track in each cat was used for final analysis due to the insufficient number of isolated single units in the L5 track of Cat A. The original, ideal design was to perform multiple mediolateral penetrations to better characterize population-level representations within a segment; however, inserting several Neuropixels probes into the narrow intradural space proved technically impractical in this preparation. In Cat B, we successfully inserted one electrode each into the L5 and L6 segments, but the L5 track yielded only a very small number of neurons that showed posture-dependent modulation and was therefore excluded from analysis. With Neuropixels 2.0 (Steinmetz *et al*., 2021) or comparable slim-shank arrays (Zhao *et al*., 2023), such dense sampling may become feasible and is a focus for future work.

Second, due to individual differences in hindlimb size, the workspace covered by the robotic arm differed between animals. Specifically, the larger body size of Cat B resulted in a smaller relative area covered by the 16 endpoint positions compared with Cat A. This difference in the spatial extent of endpoint manipulation could partially explain why Cat B showed a smaller range of firing-rate modulation and fewer neurons with statistically significant tuning. Consequently, between-animal differences (for example, a larger fraction of single–joint-angle neurons in Cat B) may reflect either individual anatomical factors or differences in electrode locus within the spinal cord.

## Acknowledgments

We would like to thank Amit Yaron for installing the high-density recording system, Moeko Kudo for her technical assistance, and Akito Kosugi and Trevor Smith for their help in constructing the analysis code. This work was partially supported by Grants-in-Aid JP18020030, JP18047027, and JP26120003 (to K.S.), and JP19H05728 and JP24K00833 (to T.F.) from the Ministry of Education, Culture, Sports, Science and Technology of Japan (MEXT). Additional support was provided in part by the NSF CRCNS Japan–US grant 2113096 to K.S. (Subaward PI) and Francisco Valero-Cuevas (PI).

## Author Contributions

K.S. and K.T. designed the study. Y.S., K.M., S.E., E.C., S.F., M.T., K.T., and K.S. performed the experiments. Y.S., K.M., and T.F. analyzed the data. Y.S., T.F., and K.S. wrote the manuscript. All authors approved the final version of the manuscript.

---

**Figure S1.**
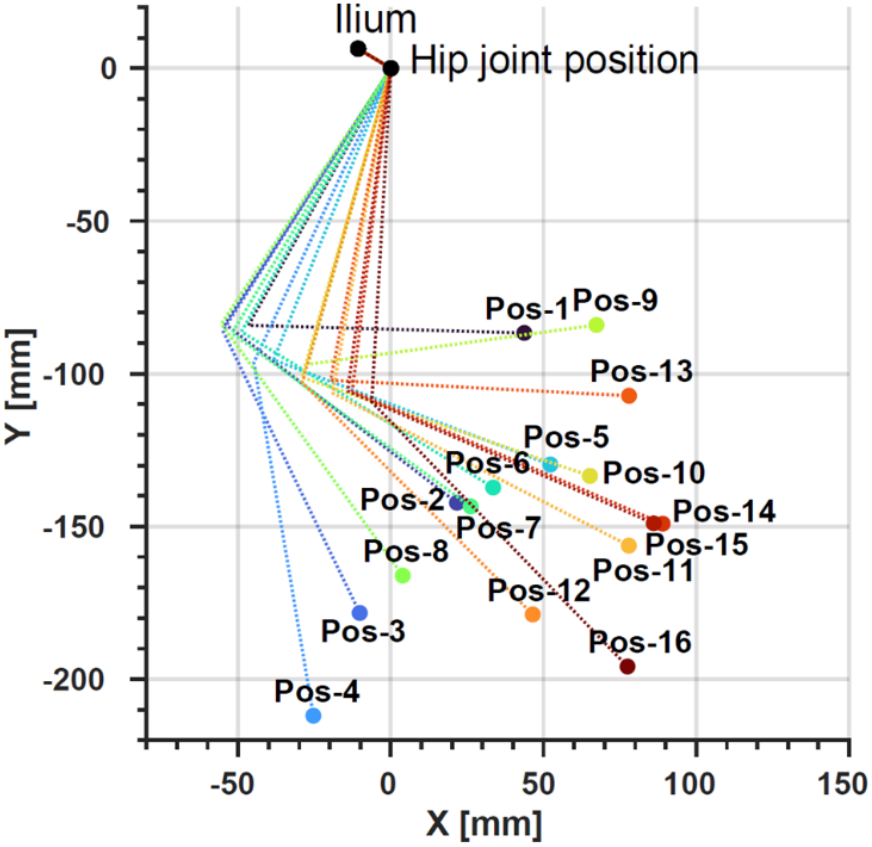
X-Y positions of the ilium, hip, knee, and ankle joints for estimating the recording position in Cat B. Data representation is the same as in Figure 2D.

**Figure S2.**
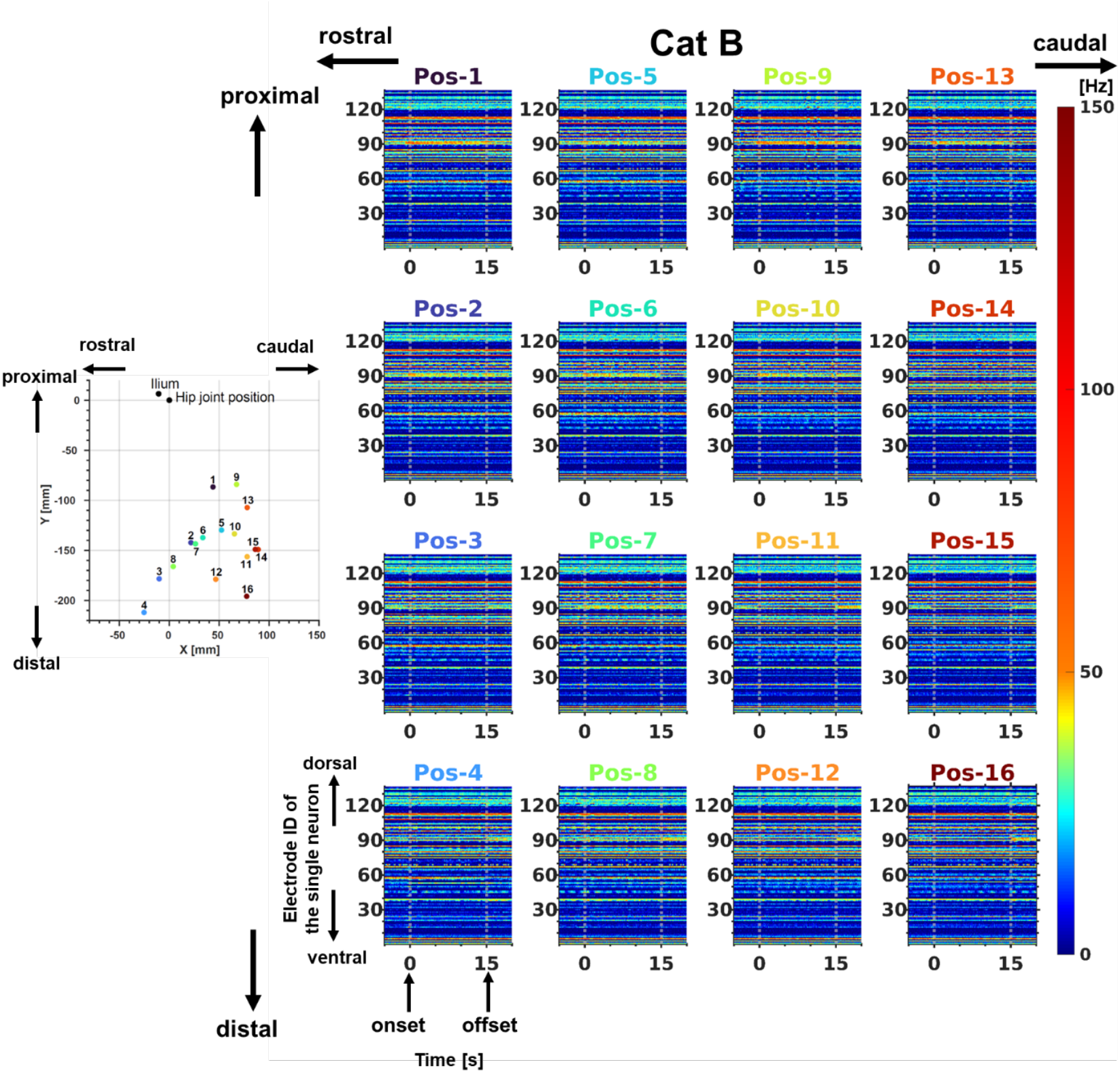
Activity of 136 single neurons across 16 recording positions in Cat B. The right panels shows the neural activity of 136 single neurons at each corresponding recording position; other details are as described in Figure 3B. The spatial arrangement of the right panels follows the configuration shown in Figure 4.

## References

Bosco G & Poppele RE. (1993). Broad directional tuning in spinal projections to the cerebellum. J Neurophysiol 70, 863–866.

Bosco G & Poppele RE. (1997). Representation of multiple kinematic parameters of the cat hindlimb in spinocerebellar activity. J Neurophysiol 78, 1421–1432.

Bosco G & Poppele RE. (2001). Proprioception from a spinocerebellar perspective. Physiol Rev 81, 539–568.

Bosco G, Poppele RE & Eian J. (2000). Reference frames for spinal proprioception: limb endpoint based or joint-level based? J Neurophysiol 83, 2931–2945.

Bosco G, Rankin A & Poppele R. (1996). Representation of passive hindlimb postures in cat spinocerebellar activity. J Neurophysiol 76, 715–726.

Brown AG & Fyffe RE. (1981). Form and function of dorsal horn neurones with axons ascending the dorsal columns in cat. J Physiol 321, 31–47.

Buford JA & Smith JL. (1990). Adaptive control for backward quadrupedal walking. II. Hindlimb muscle synergies. J Neurophysiol 64, 756–766.

Chowdhury RH, Glaser JI & Miller LE. (2020). Area 2 of primary somatosensory cortex encodes kinematics of the whole arm. Elife 9.

Clark FJ & Burgess PR. (1975). Slowly adapting receptors in cat knee joint: can they signal joint angle? J Neurophysiol 38, 1448–1463.

Degenhart AD, Bishop WE, Oby ER, Tyler-Kabara EC, Chase SM, Batista AP & Yu BM. (2020). Stabilization of a brain-computer interface via the alignment of low-dimensional spaces of neural activity. Nat Biomed Eng 4, 672–685.

Fathi Y & Erfanian A. (2022). Decoding Bilateral Hindlimb Kinematics From Cat Spinal Signals Using Three-Dimensional Convolutional Neural Network. Front Neurosci 16, 801818.

Ferrell WR. (1980). The adequacy of stretch receptors in the cat knee joint for signalling joint angle throughout a full range of movement. J Physiol 299, 85–99.

Goodman JM, Tabot GA, Lee AS, Suresh AK, Rajan AT, Hatsopoulos NG & Bensmaia S. (2019). Postural Representations of the Hand in the Primate Sensorimotor Cortex. Neuron 104, 1000–1009 e1007.

Grea H, Desmurget M & Prablanc C. (2000). Postural invariance in three-dimensional reaching and grasping movements. Exp Brain Res 134, 155–162.

Grigg P. (1975). Mechanical factors influencing response of joint afferent neurons from cat knee. J Neurophysiol 38, 1473–1484.

Grigg P & Greenspan BJ. (1977). Response of primate joint afferent neurons to mechanical stimulation of knee joint. J Neurophysiol 40, 1–8.

Hiebert GW, Whelan PJ, Prochazka A & Pearson KG. (1996). Contribution of hind limb flexor muscle afferents to the timing of phase transitions in the cat step cycle. J Neurophysiol 75, 1126–1137.

Jaeger GH, Marcellin-Little DJ, Depuy V & Lascelles BD. (2007). Validity of goniometric joint measurements in cats. Am J Vet Res 68, 822–826.

Jankowska E & Puczynska A. (2008). Interneuronal activity in reflex pathways from group II muscle afferents is monitored by dorsal spinocerebellar tract neurons in the cat. J Neurosci 28, 3615–3622.

Klaes C, Kellis S, Aflalo T, Lee B, Pejsa K, Shanfield K, Hayes-Jackson S, Aisen M, Heck C, Liu C & Andersen RA. (2015). Hand Shape Representations in the Human Posterior Parietal Cortex. J Neurosci 35, 15466–15476.

Krutki P, Jelen S & Jankowska E. (2011). Do premotor interneurons act in parallel on spinal motoneurons and on dorsal horn spinocerebellar and spinocervical tract neurons in the cat? J Neurophysiol 105, 1581–1593.

Lacquaniti F & Maioli C. (1994). Independent control of limb position and contact forces in cat posture. J Neurophysiol 72, 1476–1495.

Llobet V, Wyngaard A & Barbour B. (2022). Automatic post-processing and merging of multiple spike-sorting analyses with <em>Lussac</em>. bioRxiv, 2022.2002.2008.479192.

Maciejowski JM. (2002). Predictive Control: With Constraints. Prentice Hall.

Mathis A, Mamidanna P, Cury KM, Abe T, Murthy VN, Mathis MW & Bethge M. (2018). DeepLabCut: markerless pose estimation of user-defined body parts with deep learning. Nat Neurosci 21, 1281–1289.

Nath T, Mathis A, Chen AC, Patel A, Bethge M & Mathis MW. (2019). Using DeepLabCut for 3D markerless pose estimation across species and behaviors. Nat Protoc 14, 2152–2176.

Ostry DJ, Darainy M, Mattar AA, Wong J & Gribble PL. (2010). Somatosensory plasticity and motor learning. J Neurosci 30, 5384–5393.

Pachitariu M, Sridhar S, Pennington J & Stringer C. (2024). Spike sorting with Kilosort4. Nat Methods 21, 914–921.

Proske U & Gandevia SC. (2012). The proprioceptive senses: their roles in signaling body shape, body position and movement, and muscle force. Physiol Rev 92, 1651–1697.

Rexed B. (1952). The cytoarchitectonic organization of the spinal cord in the cat. J Comp Neurol 96, 414–495.

Rouse AG & Schieber MH. (2016). Spatiotemporal Distribution of Location and Object Effects in Primary Motor Cortex Neurons during Reach-to-Grasp. J Neurosci 36, 10640–10653.

Rule ME, Loback AR, Raman DV, Driscoll LN, Harvey CD & O’Leary T. (2020). Stable task information from an unstable neural population. Elife 9.

Rustioni A & Kaufman AB. (1977). Identification of cells or origin of non-primary afferents to the dorsal column nuclei of the cat. Exp Brain Res 27, 1–14.

Sangole AP & Levin MF. (2008). Palmar arch dynamics during reach-to-grasp tasks. Exp Brain Res 190, 443–452.

Shadmehr R & Mussa-Ivaldi FA. (1994). Adaptive representation of dynamics during learning of a motor task. J Neurosci 14, 3208–3224.

Sharma KK, Diltz MA, Lincoln T, Albuquerque ER & Romanski LM. (2024). Neuronal Population Encoding of Identity in Primate Prefrontal Cortex. J Neurosci 44.

Siegle JH, Lopez AC, Patel YA, Abramov K, Ohayon S & Voigts J. (2017). Open Ephys: an open-source, plugin-based platform for multichannel electrophysiology. J Neural Eng 14, 045003.

Soechting JF & Flanders M. (1992). Moving in three-dimensional space: frames of reference, vectors, and coordinate systems. Annu Rev Neurosci 15, 167–191.

Standardization. IOf. (2025). Robotics - Safety requirements Part 1: Industrial robots. Geneva.

Stein RB, Weber DJ, Aoyagi Y, Prochazka A, Wagenaar JB, Shoham S & Normann RA. (2004). Coding of position by simultaneously recorded sensory neurones in the cat dorsal root ganglion. J Physiol 560, 883–896.

Steinmetz NA, Aydin C, Lebedeva A, Okun M, Pachitariu M, Bauza M, Beau M, Bhagat J, Bohm C, Broux M, Chen S, Colonell J, Gardner RJ, Karsh B, Kloosterman F, Kostadinov D, Mora-Lopez C, O’Callaghan J, Park J, Putzeys J, Sauerbrei B, van Daal Rjj, Vollan AZ, Wang S, Welkenhuysen M, Ye Z, Dudman JT, Dutta B, Hantman AW, Harris KD, Lee AK, Moser EI, O’Keefe J, Renart A, Svoboda K, Hausser M, Haesler S, Carandini M & Harris TD. (2021). Neuropixels 2.0: A miniaturized high-density probe for stable, long-term brain recordings. Science 372.

Tanaka Y & Hirai N. (1994). Physiological studies of thoracic spinocerebellar tract neurons in relation to respiratory movement. Neurosci Res 19, 317–326.

Tibshirani R. (1996). Regression Shrinkage and Selection Via the Lasso. Journal of the Royal Statistical Society: Series B (Methodological) 58, 267–288.

Tsay JS, Kim H, Haith AM & Ivry RB. (2022). Understanding implicit sensorimotor adaptation as a process of proprioceptive re-alignment. Elife 11.

Vallbo AB. (1974). Afferent discharge from human muscle spindles in non-contracting muscles. Steady state impulse frequency as a function of joint angle. Acta Physiol Scand 90, 303–318.

Veshchitskii A, Shkorbatova P & Merkulyeva N. (2022). Neurochemical atlas of the cat spinal cord. Front Neuroanat 16, 1034395.

Wei JY, Simon J, Randic M & Burgess PR. (1986). Joint angle signaling by muscle spindle receptors. Brain Res 370, 108–118.

Yaguchi H, Takei T, Kowalski D, Suzuki T, Mabuchi K & Seki K. (2015). Modulation of spinal motor output by initial arm postures in anesthetized monkeys. J Neurosci 35, 6937–6945.

Yakovenko S, McCrea DA, Stecina K & Prochazka A. (2005). Control of locomotor cycle durations. J Neurophysiol 94, 1057–1065.

Zhao Z, Zhu H, Li X, Sun L, He F, Chung JE, Liu DF, Frank L, Luan L & Xie C. (2023). Ultraflexible electrode arrays for months-long high-density electrophysiological mapping of thousands of neurons in rodents. Nat Biomed Eng 7, 520–532.

